# *Gn1a* rice promotes symbiotic fish growth via reprogramming soil microbiome

**DOI:** 10.64898/2026.01.13.699285

**Authors:** Shen-Zheng Zeng, Yu-Chan Zhang, Rui-Rui He, Hui-Yin Pang, Zhi-Jian Huang, Zhi-Xuan Deng, Ren-Jun Zhou, Xiao-Cheng Tang, Tian-Hao Li, Juan-Rong Lv, Qiao-Juan Huang, Jia-Biao Yang, Shao-Ping Weng, Jian-Guo He, Yue-Qin Chen

## Abstract

Rice-fish symbiotic ecosystems significantly contribute to world food supply. Mutation in rice (*Oryza sativa*) quantitative trait locus *Gn1a*, encoding a cytokinin oxidase/dehydrogenase (OsCKX2), leads to cytokinin accumulation in plant, which promotes tillering and panicle branching to achieved higher yield and also shows a selective effect on soil microbial composition. Given that the growth of aquatic animals can be shaped by their surrounding microbial communities, we asked whether adjusting rice cytokinin levels might synergistically boosts both rice yield and fish growth. Herein, we co-cultured common carp (*Cyprinus carpio*) with *ckx2* rice plants to determine the effect of cytokinins on fish. Surprisingly, compared to wild-type plants, the *ckx2* plants enhanced nearly 10% fish growth and improved fish antibacterial capacity against *Aeromonas hydrophila* and *Edwardsiella tarda*. In fact, the growth-promoting effect on fish requires mediation by the soil microbiota. Mechanistically, *ckx2* plants secreted more cytokinins into soil to reprogram the microbial composition, which subsequently translocated to fish gut and modulated the keystone microbes in gut to facilitate short-chain fatty acid production and carbohydrate degradation. These findings provide the first evidence that rice-derived cytokinins exert a cross-kingdom growth-promoting effect on fish, providing a boosting dual-production strategy for advancing rice-fish symbiotic ecosystem.

## Introduction

Rice (*Oryza sativa*) is a staple crop for over half of the world’s population and a cornerstone of global food security^1^. Rice-fish symbiotic culture systems, including rice-fish, rice-crayfish, rice-shrimp and rice-crab co-culture, foster mutualistic interactions among diverse biological components to acquire more food, high-quality protein and economic income^2–6^. Compared to rice-only farming, rice-fish coculture could increase 25% of more food (rice and fish) production, and raise economic water productivity from US$0.19/m^3^ in rice monoculture to US$0.41/m^3 7^, leading to their implementation in more than 30 countries across Asia, Africa and Americas, particularly among small-scale farmers in rural regions^8–10^. Within these symbiotic systems, fish predation suppresses rice pests and *Daphnia*, and drives the cycling of nitrogen and phosphorus, helping lower pesticide by >60% and fertilizer uses by >20%^11–13^. Meanwhile, rice plants are reported to improve the quality of co-cultured fish, including tilapia (*Oreochromis niloticus*) and common carp (*Cyprinus carpio*), with higher muscle protein and fat content, higher gut microbial diversity, and distinctive gut and liver metabolite profiles^5,14^.

Cultivated rice varieties underwent substantial improvements in grain yield and quality during domestication. Cytokinin phytohormones are synthesized mainly in plant root tips and serve as pivotal regulators of plant architecture, coordinating growth and development across diverse organs, such as tillering and panicle branching^15–17^, which thus regulates rice grain yield^18,19^. Importantly, considerable concentrations of various cytokinins are also detected in the soil^20^. Appropriate cytokinin accumulation is considered an indicator of healthy soil^21^, and has selective effects on soil and phyllosphere microbial composition^22,23^. Cytokinins promote the abundance of specific microbes, including Bacilli and Bacteroidia^24–26^. These two taxa are often abundant in the gut of aquatic animals and are widely used as probiotics because they promote immunity and growth^27–31^. Given that the gut microbiomes of aquatic animals can be shaped by their surrounding microbial communities^32–35^, we asked whether, in a rice-fish eco-symbiotic culture system, adjusting rice cytokinin levels might synergistically boosts both rice yield and fish growth via reprogramming of the fish microbiota. Answering this question could be central to unlocking the full potential of eco-symbiotic culture systems.

The quantitative trait locus *Gn1a* encoded a cytokinin oxidase/dehydrogenase (OsCKX2). In the present study, we used a rice mutant with knockout of *OsCKX2* that catalyzes the degradation of active cytokinins, which leads to cytokinin accumulation in inflorescence meristems, contributing to nearly 16% more grains per panicle and 17% higher grain yield per plant^36^. We then established a controlled eco-symbiotic system by co-cultivating wild-type (WT) or *ckx2* plants with common carp (*Cyprinus carpio*). Strikingly, the mutant plants enhanced fish growth and antibacterial capacity in both laboratory and field experiments. Mechanistically, loss of *OsCKX2* promotes cytokinin secretion into the soil, which reshapes the soil microbiota and alters keystone taxa in the fish gut microbiome. This study is the first to report the mechanisms that *ckx2* plants secrete cytokinins to enhance fish growth and immunity in the rice-fish co-culture system, highlighting promising applications for sustainable integrated aquaculture.

## Results

### *OsCKX2* loss of function plants promote fish growth in a symbiotic culture system

Since loss function of *OsCKX2* promote yield production due to the accumulation of cytokinins^37^, we asked whether, in a rice-fish eco-symbiotic culture system, applying this high-yield rice variant also exert an effect on fish growth? To address this question, we established a controlled in-house symbiotic system by co-cultivating *ckx2* or wild-type plants (WT) plants with common carp ((*Cyprinus carpio*, 3 tanks for each group), growing rice plants from the seedling stage to harvest in a tank with soil and introducing fish (*n* = 60, 9.55±1.47g) at 28 days after rice germination (**Fig. 1a**). Fish was collected at seven time-points (0, 15, 30, 40, 50, 60 and 70 days after co-cultivation, DAC) to measure their body weight and length.

**Fig. 1.**
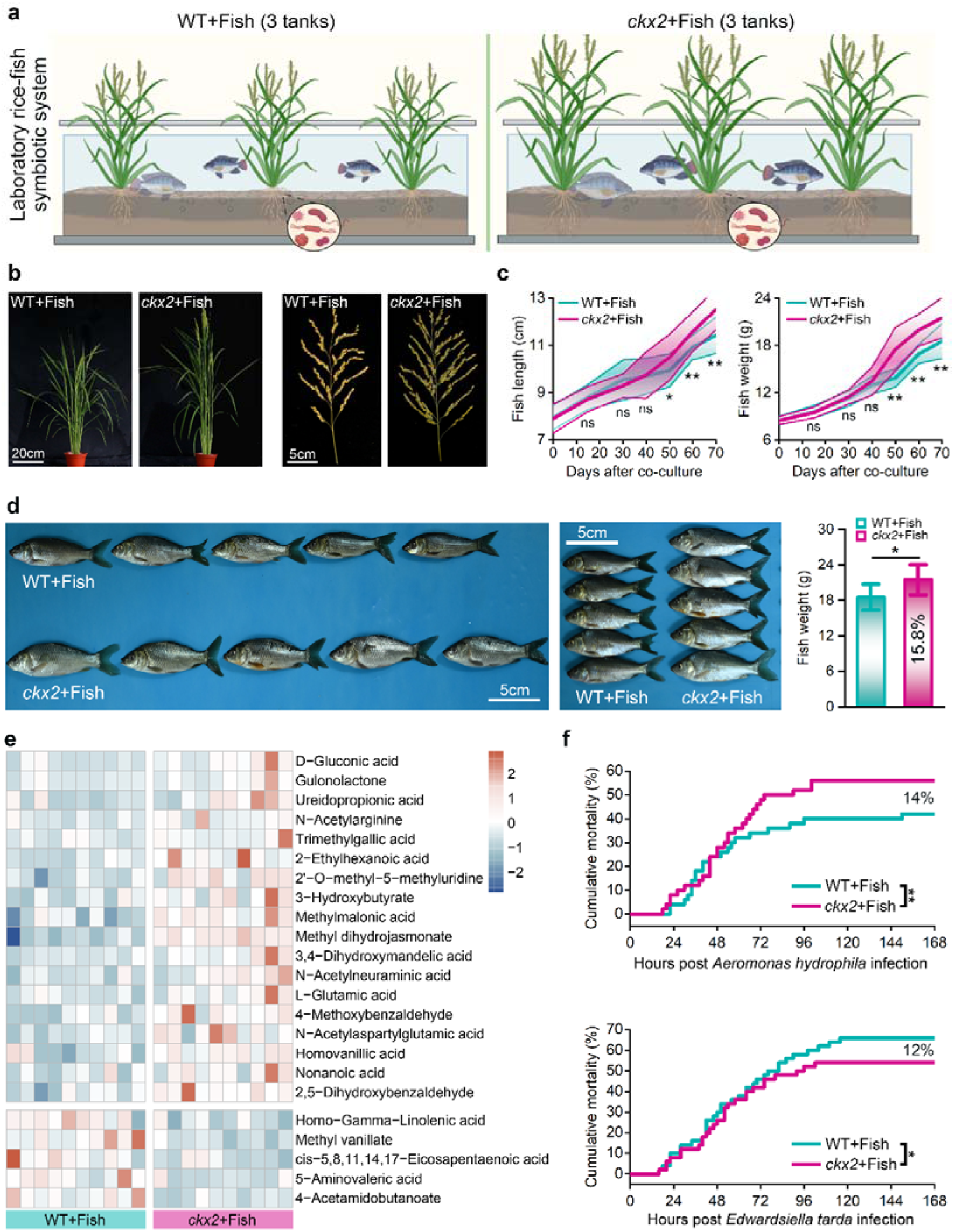
*ckx2* rice plants promotes both grain yield and fish growth in rice-fish symbiotic ecosystem in laboratory. **(a)** Diagram illustrating the experimental co-culture system. Common carp individuals were divided into two groups and co-cultured with either WT rice plants or with *ckx2* plants, with three independent replicates per group. **(b)** Representative photographs of rice plants showing yield-related traits, assessed at 70 days after co-cultivation (DAC). Scale bars, 20 cm (whole plants), 5 cm (panicles). **(c)** Quantification of fish length (left) and fish weight (right) at 15, 30, 40, 50, 60 and 70 DAC (*n* = 5 per replicate). Asterisks indicate significant differences, based on a two-tailed *t*-test; * *P* < 0.05; ** *P* < 0.01; ns, not significant (for each group, fish: *n* = 15). **(d)** Representative photographs of fish co-cultured with WT or *ckx* plants at 70 DAC. Fish co-cultured with *ckx2* plants were 15.8% heavier than controls. Scale bars, 5 cm. Asterisks indicate significant differences, based on a two-tailed *t*-test; * *P* < 0.05. **(e)** Heatmap representation of compound contents in the gut of fish co-cultured with WT or *ckx2* plants, as determined by high-performance liquid chromatography. A total of 23 compound types of fatty acids and lipids with significantly different contents between the two groups were identified (*p* < 0.05, two-tailed *t*-test; for each group, fish: *n* = 10). The concentration of these compounds was normalized by Z-score in the heatmap. **(f)** Assessment of bacterial resistance of carp infected with *Aeromonas hydrophila* (top) or *Edwardsiella tarda* (bottom) and co-cultured with WT or *ckx2* plants (for each group, fish: *n* = 50). Significant differences between the two groups were assessed by the log-rank test; * *P* < 0.05; ** *P* < 0.01.

Consistent with observations under field conditions, the *ckx2* plants produced larger panicles and more seeds per plant than WT plants, suggesting the stability of its desirable agronomic traits in the symbiotic system, (**Fig. 1b**). Very unexpected, we observed that fish co-cultured with *ckx2* plants exhibited significantly superior growth, surpassing those co-cultured with WT rice in both weight and length at 50 DAC and increasing progressively over time (**Fig. 1c**). By 70 DAC, the fish growth co-cultured with *ckx2* plants showed an increase of 15.8% in average body weight compared with those co-cultured with WT plants (**Fig. 1d**).

Interested by this growth boost, we further measured the fish metabolic and immune traits. The *ckx2*-co-cultured fish displayed elevated muscle fat content (28.1%), heightened activities of amylase (25.4%) and lipase (10.1%, **Fig. S1**), and a 11.4% rise in total lipid in fish gut, compared to those co-cultured with WT plants (**Fig. 1e, Fig. S2**). Notably, we detected an upregulation of specific metabolites like D-gluconic acid and 3-Hydroxybutyrate in fish gut, which are reported to improve resistance to bacterial pathogens in animals ^38^. This metabolic shift was paralleled with the significant upregulation of antibacterial genes in fish (**Fig. S3**), suggesting a fortified immune state in fish co-cultured with *ckx2* plants. To confirm this observation, we performed an antibacterial challenge. When infected with *Aeromonas hydrophila* or *Edwardsiella tarda*, fish co-cultured with *ckx2* plants showed a significantly lower mortality than that of control (56% to 42%, 66% to 54% respectively; **Fig. 1f**). Taken together, these results demonstrate that genetically inactivating *OsCKX2* enhanced grain yield and conferred dual benefits to fish by promoting growth and strengthening antimicrobial capacity in a symbiotic ecosystem.

### Secretion of cytokinins from *ckx2* plants into the symbiotic co-culture system improves fish growth

The above observations showed that knocking out *OsCKX2* enhanced rice grain yield and promoted the growth and quality of co-cultivated fish. Cytokinins are synthesized mainly in plant root tips and serve as long-distance signaling molecules, coordinating growth and development across diverse organs^39–41^. Given that cytokinin accumulation is the hallmark feature resulting from the *OsCKX2* knockout, we therefore asked whether *ckx2* plants promote fish growth via secreting cytokinins into the symbiotic environment. To address this question, we co-cultivated rice plants (WT or *ckx2*) and common carp in a paddy-pond co-culture system with rice plants placed around a central fishpond, using an exogenous addition of 3 μg/L synthetic cytokinin 6-benzylaminopurine (6-BA) to WT rice for comparison (**Fig. 2a**).

**Fig. 2.**
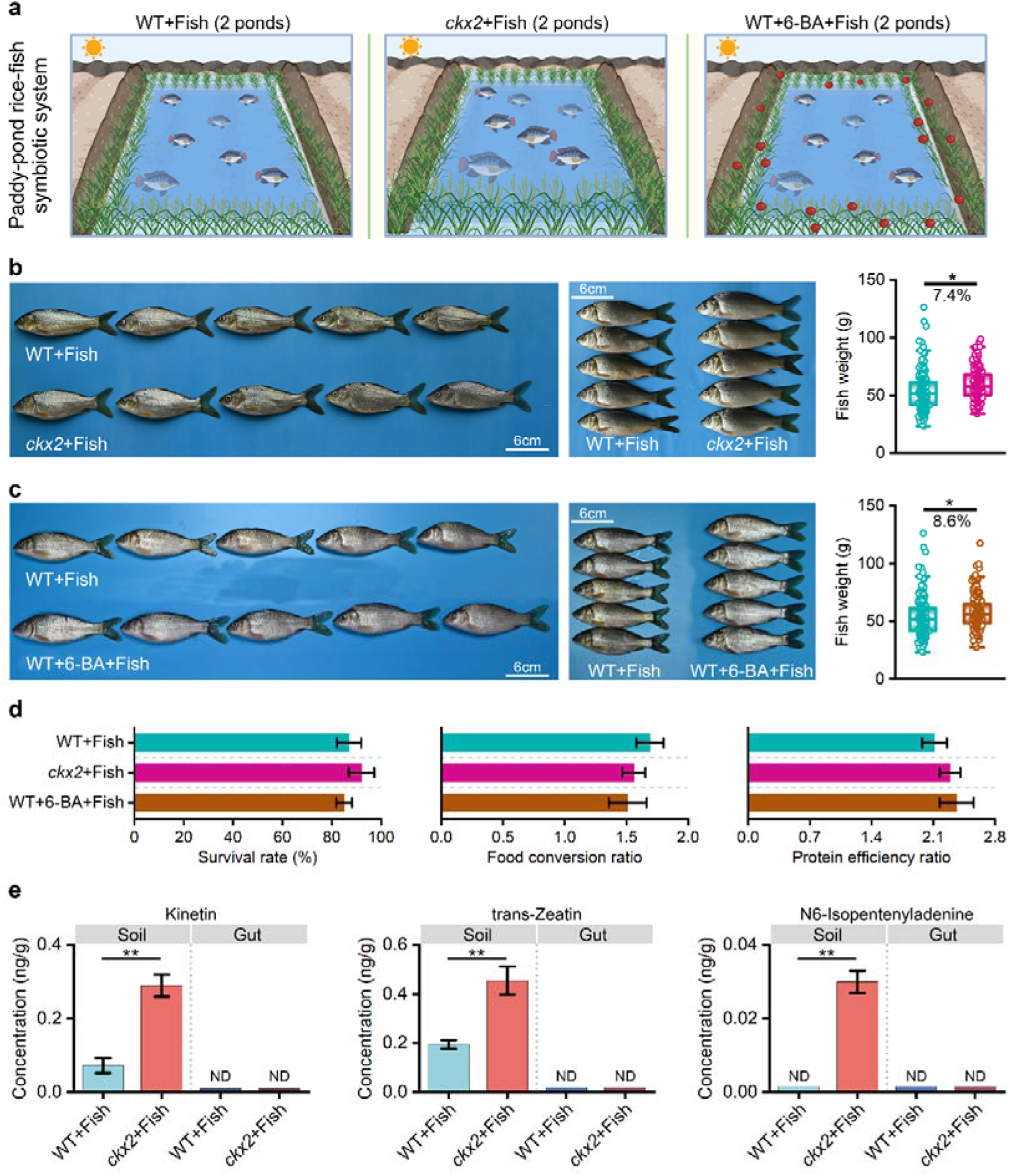
Paddy-pond symbiotic system confirms that *ckx2* plants enhance fish growth via secreting cytokinins into the co-culture system. (a) Diagram of the paddy-pond co-culture system with WT or *ckx2* plants (6 m × 9 m; *n* = two ponds) cultivated around a central fishpond (4 m × 6 m) with 300 carp. The experiment was conducted twice, in the autumn of 2024 and the spring of 2025. Fish was co-cultured with rice plants (WT or *ckx2*). To evaluate the effect of cytokinin on fish growth, an exogenous addition of 3 μg/L synthetic cytokinin 6-benzylaminopurine (6-BA) to was supplemented to WT rice weekly for comparison. **(b)** Representative photographs (left) and quantification of weight (right) from fish co-cultured with WT or *ckx2* plants at 70 DAC. Scale bars, 6 cm. The asterisk indicates significant differences, based on a two-tailed *t*-test; * *P* < 0.05 (for each group, fish: *n* = 400). Data from the additional trial (Spring 2025), presented in Fig. S4, also demonstrate a significant increase in body weight in the *ckx2*+Fish group. **(c)** Representative photographs (left) and quantification of weight (right) from fish co-cultured with WT plants alone or with cytokinin treatment at 70 DAC. Scale bars, 6 cm. The asterisk indicates significant differences, based on a two-tailed *t*-test; \**P* < 0.05 (for each group, fish: *n* = 400). Data from the additional trial (Spring 2025), presented in Fig. S4, also demonstrate a significant increase in body weight in the WT+6-BA+Fish. **(d)** Survival rate, feed conversion ratio and protein efficiency ratio of each pond at 70 DAC (for each group, pond: *n =* 2). **(e)** Detection of cytokinin contents in soil and fish gut using high-performance liquid chromatography. Data are presented as mean ± standard deviation (for each group, soil: *n* = 4; gut: *n* = 4). ND represents the not detected. Significant differences between the two groups were assessed by t-test; \*\**P* < 0.01.

Consistent with observations in laboratory experiment (Fig. 1C), fish co-cultivated with *ckx2* plants grew faster than those co-cultivated with WT plants, ending 7.4% heavier than the WT+Fish group at 70 DAC (**Fig. 2b**, **Fig. S4**). More importantly, we observed that adding cytokinin to WT plants pond similarly boosted fish weight by 8.6% (**Fig. 2c**), along with a better feed conversion ratio and protein efficiency ratio for fish production (**Fig. 2d**).

To investigate whether higher concentration of cytokinin accumulated in the *ckx2* plants contributes to the fish growth, we conducted a liquid chromatography-tandem mass spectrometry to quantify different cytokinin in both soil and fish gut samples. A significant elevation of three cytokinin variants were detected in the soil around *ckx2* plant roots, including kinetin (0.073 to 0.29 ng/g), trans-Zeatin (0.19 to 0.43 ng/g) and N6-isopentenyladenine (<0.01 to 0.03 ng/g, **Fig. 2e**). However, none of these cytokinins were detected in the fish guts (**Fig. 2e**). These results suggested that *ckx2* plant secreting cytokinins could improve fish growth, yet cytokinins did not directly accumulate in fish, indicating an indirect mechanism for the growth promotion.

### Cytokinins reprogram the microbial composition of soil and fish gut

Given the profound influence of the gut microbiota on fish growth, and cytokinins exert selective pressure on specific microbes and even the overall microbial structure^24–26^, we next examined whether cytokinins reprogram the microbial compositions of soil and fish gut. To reveal this ecological cascade, we collected soil and gut samples from co-culture systems with WT or *ckx2* plants at seven time-points (as in Fig. 1C), which were subjected to 16S rDNA amplicon sequencing for microbiome profiling. After quality control, 10,346 amplicon sequence variants (ASVs) in soil samples and 6,198 ASVs in fish gut samples were obtained (**Fig. S5**).

A temporal dynamic emerged that the soil microbial structure in *ckx2*+Fish group began to cluster apart from those in WT+Fish group at 30 DAC (*P* = 0.030), while significant structural shift in fish gut became apparent from 50 DAC (*P* = 0.024, **Fig. 3a**). Over time, the influence of *ckx2* plants amplified, accounting for substantial variance in the community structures (soil: *R*^2^ = 0.275; gut: *R*^2^ = 0.391, **Fig. 3b**). Interestingly, the *ckx2* plants did not significantly affect the alpha diversity of soil bacterial communities, but were associated with significantly lower microbial diversity in the gut microbiome at later stages (60 and 70 DAC, **Fig. 3c**).

**Fig. 3.**
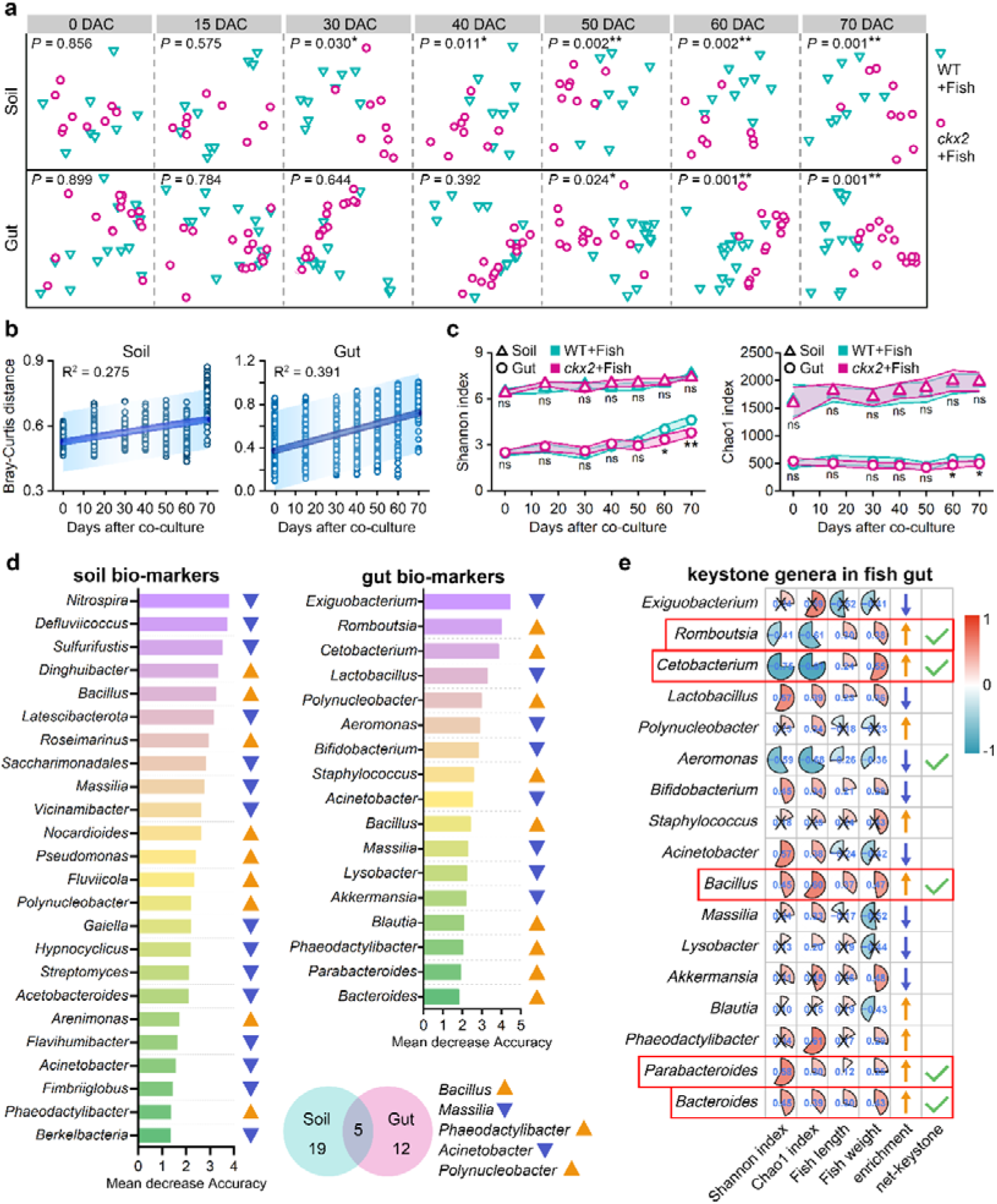
*ckx2* plants alter the microbial composition in the soil and fish gut. **(a)** At each timepoint, three soil sample and five gut samples were collected at each tank. Principal coordinate analysis based on Bray-Curtis distance was applied to evaluate the difference in microbial structure between WT+Fish and *ckx2*+Fish groups. Green triangles indicate the WT+Fish group, and purple circles indicate the *ckx2*+Fish group (for each group, soil: *n* = 9; gut: *n* = 15). Significant difference (*P* < 0.05) in microbial structure between groups was assessed by permutational multivariate analysis of variance (PerMANOVA). **(b)** Temporal dynamics of structural divergence between WT+Fish and *ckx2*+Fish groups was assessed based on Bray-Curtis distance. The blue lines represent the least-squares regression fits, and the shaded area represents the 95% confidence intervals. The coefficient of determination (*R*^2^) for the least-squares regression fit is shown. **(c)** Comparison of alpha diversity between WT+Fish and *ckx2*+Fish groups using Shannon index (left) and Chao index (right) (for each group, soil: *n* = 9; gut: *n* = 15). Significant differences between the two groups were assessed by the log-rank test; * *P* < 0.05; ** *P* < 0.01; ns, no significance. **(d)** Identification of soil and gut microbial biomarkers for *ckx2*+Fish group based on a random-forest model. Biomarkers were sorted by their mean decrease accuracy in the random-forest model. Blue triangles represent taxa with increased abundance in *ckx2*+Fish group, whereas orange triangles represent those with decreased abundance. The inset Venn diagram shows the extent of overlap between marker genera from the soil and fish gut samples. **(e)** Identification of keystone microbes in fish gut by examining their correlations with alpha diversity, body weight, and length, their response to *ckx2* plants, and their topological roles within the microbial co-occurrence network. The cross represents no significance (*P* > 0.05) in Spearman’s rank correlation, and the size of sector represents the *r* value. The five genera highlighted by red boxes were identified as the keystone microbes based on the aforementioned criteria.

To identify the bacterial taxa responsive to cytokinins and the keystone microbes in the fish gut (defined as hub-nodes within the species interaction network and linked to community diversity). We identified 21 genera in soil and 17 in fish gut as biomarkers for the *ckx2* plants group (**Fig. 3d, Fig. S6**). These biomarkers effectively distinguished soil and gut microbiomes between WT +Fish and *ckx2*+Fish groups with high accuracy (60.0-88.9%, **Fig. S7**). Five genera biomarkers, *Bacillus*, *Massilia*, *Phaeodactylibacter*, *Acinetobacter* and *Polynucleobacter*, overlapped between the soil and fish gut samples (**Fig. 3d**). Furthermore, *Cetobacterium*, *Romboutsia*, *Parabacteroides*, *Bacteroides* and *Bacillus* were identified as the keystone microbes, based on their higher abundance in the *ckx2*+Fish group, which was also supported by their positions as central hubs in the gut microbial network (**Supplementary Figures 8 and 9**), and their abundance was significantly correlated with alpha diversity, fish weight and fish length (**Fig. 3e**). These changes in microbial composition also altered the potential functions of gut microbiome, which were functionally associated with butyric acid metabolism, biosynthesis of unsaturated fatty acids, and arginine, proline and histidine metabolism (**Fig. S10**).

Notably, exogenous cytokinin to WT plants ecosystem successfully reproduced the key phenomena associated with *ckx2* plants, including higher alpha diversity and elevated abundance of keystone microbes in fish gut microbiome, along with the alteration of soil biomarkers including *Bacillus*, *Acinetobacter* and *Phaeodactylibacter* (**Fig. S11**). These results suggested that cytokinins affected the composition of the soil microbiomes, leading to the rise of keystone genera in fish gut, which might contribute to better fish growth.

### Soil microbes reprogrammed by *ckx2* plants translocate and colonize in fish gut

Having established that altered microbial composition in both soil and gut, yet cytokinins were undetectable in fish gut. We therefore established a controlled aquaculture experiments to determine whether accelerated fish growth was directly induced by cytokinins or was mediated through the soil microbial intermediaries. Following experiments were designed: (1) a full rice-soil-fish system, (2) a soil-fish system, and (3) a fish-only system. The experiments were conducted with or without cytokinin supplementation (**Fig. 4a**).

**Fig. 4.**
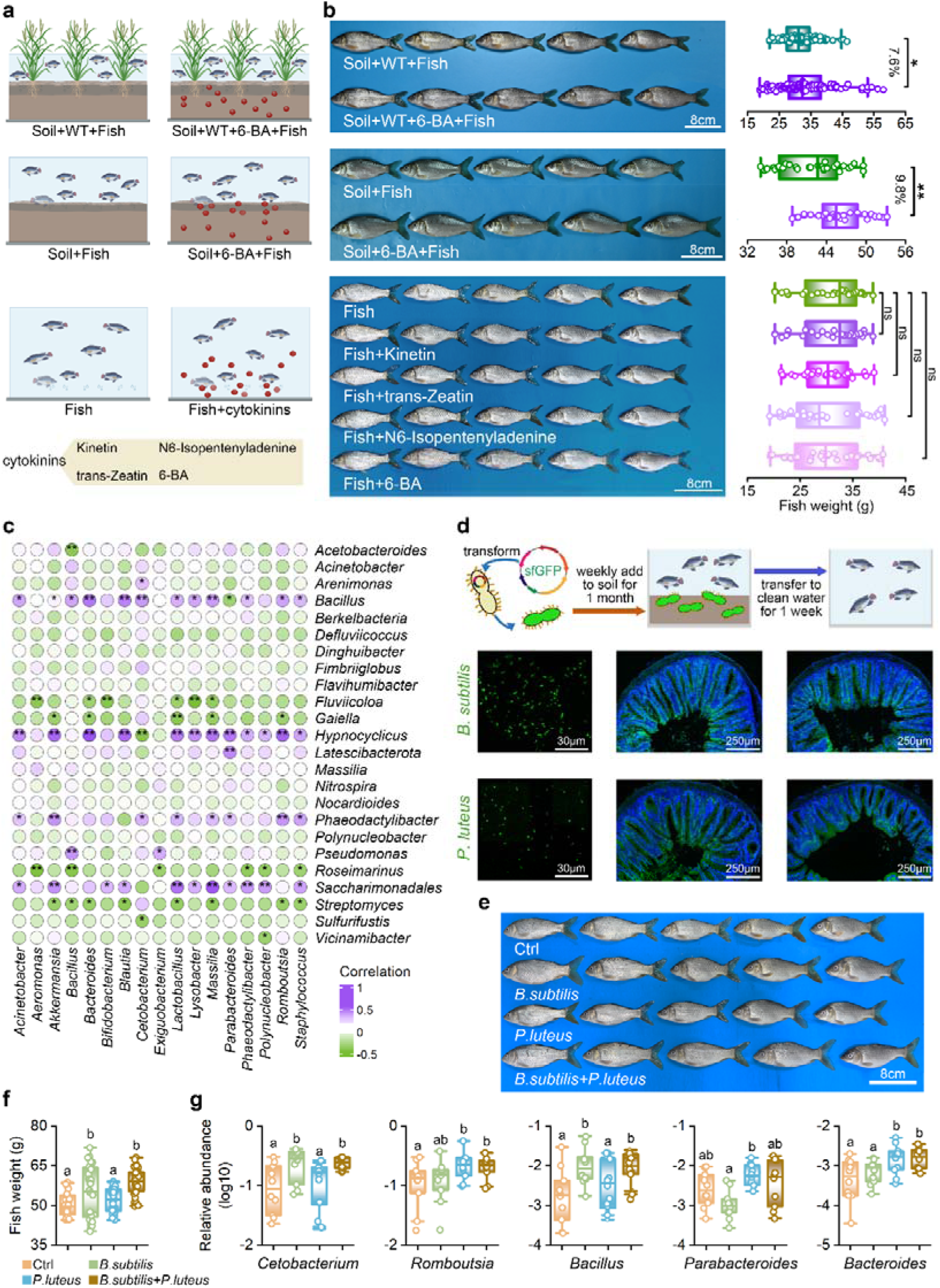
The soil microbiota serves as an intermediary between cytokinins and fish gut microbial community. **(a)** Experimental design to test whether cytokinins directly modulate the fish gut microbiota or indirectly influence microbial communities through soil microbial intermediaries. Six sets of fish were considered, excluding rice plants, soil and cytokinin treatment from the symbiotic ecosystem. Fish growth was measured at 70 DAC. **(b)** Representative photographs (left) and weight (right) of fish co-cultured with WT plants in soil or treated with cytokinins. Scale bar: 8cm. Asterisks indicate significant differences, based on a two-tailed t-test; * *P* < 0.05; ** *P* < 0.01; ns, not significant. **(c)** Spearman’s rank correlation coefficient analysis for soil and gut biomarkers. Asterisks indicate a significant correlation between two genera (\**P* < 0.05; \*\**P* < 0.01). The value of the Spearman’s rank correlation coefficient is indicated by different colors from green for negative values to purple for positive values. **(d)** Representative fluorescence micrographs of two keystone genera labeled with the green fluorescent protein (GFP). The GFP-labeled strains were weekly added to soil for 1 month, and green fluorescence was observed on the gut microvilli of fish. The green signal remained detectable one-week post-transfer to clean water following a “clearing procedure”, which proved that soil microbes could translocate and colonize in fish gut. Scale bars represent 30μm for the bacterial image and 250μm for the gut image, respectively. **(e)** Representative photograph of carp cultured with weekly supplement of B. subtilis and/or P. luteus for 70 days. Scale bar: 8cm. The experiment was conducted twice with similar results. Scale bar, 8 cm. **(f)** Comparison of fish weight with supplements of B. subtilis and/or P. luteus. Different lowercase letters indicate significant differences, as determined by a one-way analysis of variance (for each group, fish: *n* = 30). **(g)** Changes in the abundance of keystone genera in the fish gut with different supplements. Different lowercase letters indicate significant differences, as determined by a one-way analysis of variance (for each group, fish: n = 10).

As shown in **Fig. 4b**, Cytokinin supplementation significantly boosted fish growth by 7.6-9.8% in rice-soil-fish and soil-fish systems containing soil, whereas no significant difference in fish growth was observed in the fish-only system, where cytokinins applied directly to the water failed to alter fish growth (**Fig. 4b**). In addition, cytokinin treatment consistently reshaped the soil microbial structure and altered the abundance of some key genera, such as *Bacillus* and *Phaeodactylibacter* (**Fig. S12**). Similarly, these changes were faithfully mirrored in the fish gut microbiome (**Fig. S13**), aligning with the altered microbiome compositions in the *ckx2* co-culture system (Fig. 2), but again, only in the presence of soil. These results suggested that cytokinins promoted fish growth via reprogramming the soil microbiota.

Subsequently, we quantified the relative contribution of soil microbial community to fish gut microbiome. A great number of shared ASVs between soil and gut microbiome was detected, accounting for more than 45% of the total abundance of fish gut (**Fig. S14**). Significant correlation between changes in the microbial structure of gut and soil microbiota was observed, and soil microbes might be a major source for gut microbiome (**Fig. S15**). The strong correlation between keystone taxa across soil and gut was noticed (**Fig. 4c**), especially *Bacillus* and *Phaeodactylibacter,* previously identified as cytokinin-responsive microbes in soil, emerged as prime suspects for making the migration from soil to gut to alter gut microbiome.

To evaluate the effect of soil microbes on the fish, we successfully isolated two bacteria strains from soil that were linked to fish microbial variations: a *Bacillus subtilis* strain and a *Phaeodactylibacter luteus* strain. We then genetic engineered these two strains with green fluorescent protein to track their migration within the fish-rice system (**Fig. S16**). GFP-labeled strains were introduced weekly to the symbiotic system, and they were subsequently detected in fish gut, demonstrating their direct translocation capability from environment to gut (**Fig. 4d**). After 70 days of supplementation, fish weight in the sole *Bacillus subtilis* and the mixed supplementary groups exhibited significant increases of 8.0% and 11.2% in fish weight compared to control group, respectively, whereas the sole supplementary of *Phaeodactylibacter luteus* did not result in a significant increase (**Fig. 4e**). The rapid growth was accompanied by an enrichment of keystone genera in fish gut (**Fig. 4f**), which was consistent with the variations led by symbiosis with *ckx2* plants. These results demonstrated soil bacteria could transfer to fish gut, affecting the composition of gut microbiome and enhancing fish growth.

### Cytokinins-induced keystone microbes enhance short-chain fatty acids production and macromolecule carbohydrate degradation

Having identified a consortium of cytokinin-induced keystone microbes in fish gut, we sought to evaluate their benefits to fish growth. We isolated seven pivotal bacterial strains from fish gut, including *Cetobacterium somerae, Romboutsia lituseburensis*, *Parabacteroides goldsteinii*, *Parabacteroides johnsonii*, *Bacillus licheniformis*, *Bacillus aerium* and *Bacteroides xylanisolvens*, and subjected them to whole-genome sequencing (**Table. S1, Fig. S17**). The genomic blueprint was striking that each strain independently possessed a nearly complete genetic arsenal for short-chain fatty acid biosynthesis and degradation of complex carbohydrates like starch and cellulose (**Fig. 5a**). To confirm their ability to synthesize short chain fatty acids, we then inoculated fish feed with each bacterial isolate for a 48-hour fermentation period. Compared to unfermented feed, a nearly 2-fold increase in short-chain fatty acids was detected after fermentation, along with the elevation of multiple organic acids, including citric, fumaric and succinic acids (**Fig. 5b**). *C. somerae*, *R*. *lituseburensis* and *B. licheniformis* produced more acetic, propionic and butyric acids, while *P*. *johnsonii* and *B. xylanisolvens* synthesized more isovaleric, caproic and isocaproic acids. These results suggested that the cytokinin-induced keystone microbes in fish gut might utilize cellulose and starch in feed to produce short-chain fatty acids, and increase the content of small molecule carbohydrate.

**Fig. 5.**
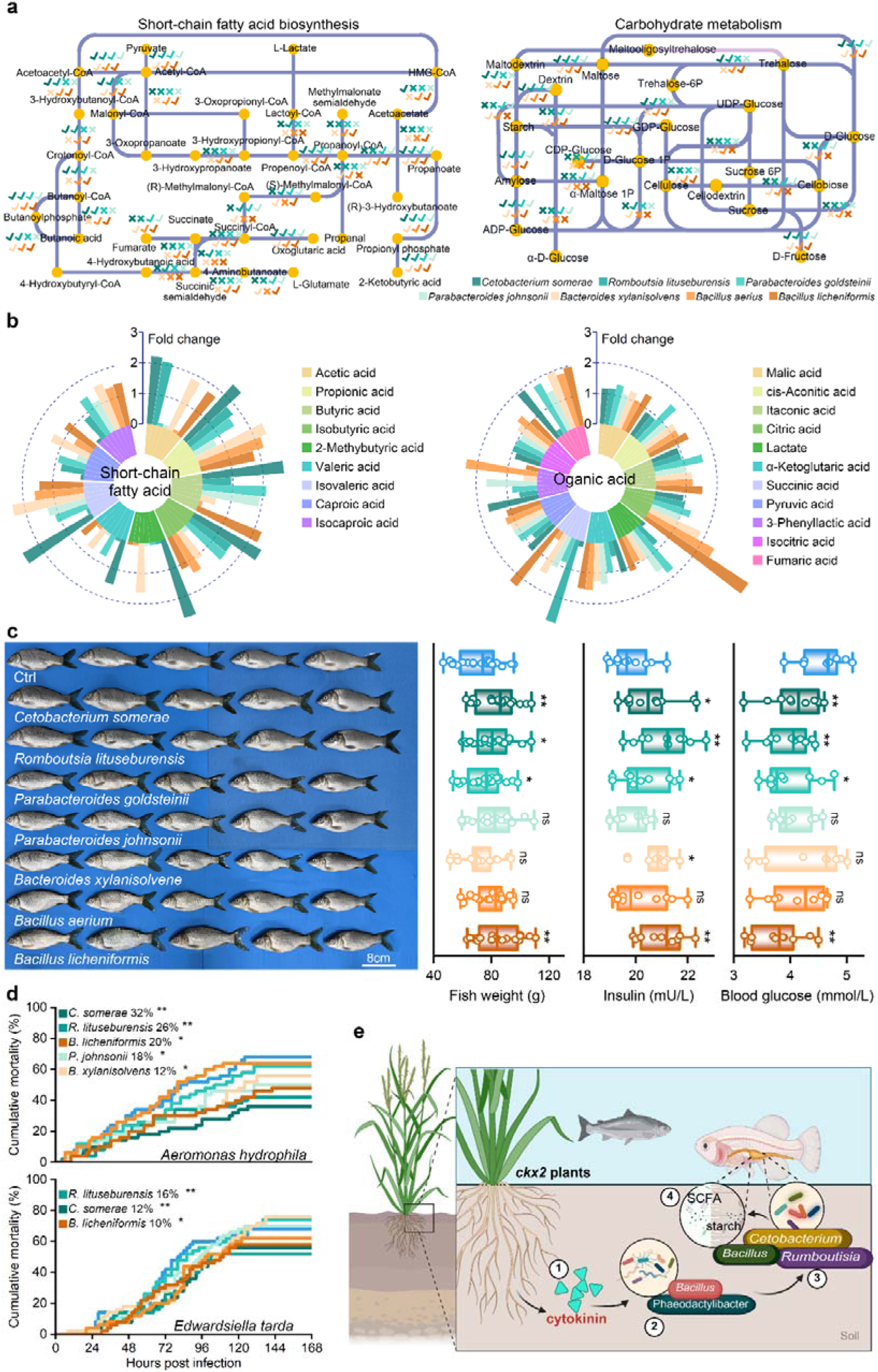
Keystone genera produce short-chain fatty acids and carbohydrate degradation to improve fish growth. **(a)** Genomic sequencing revealed that the seven strains possess functional capabilities for short-chain fatty acid biosynthesis and carbohydrate degradation. Each strain harbored a nearly complete set of genes essential for SCFA production, and showed the ability to depolymerize macromolecular substrates, such as starch and cellulose, into small molecular sugars. A hook symbol denotes the bacterium’s capability to catalyze the conversion of compounds, whereas a cross indicates the absence of this function. **(b)** A radial bar chart displayed the fold change of compounds produced after a 48-hour bacterial fermentation, compared to control group (normal feed). The compounds were categorized into two classes: short-chain fatty acids and organic acids generated from macromolecular carbohydrate degradation. **(c)** Representative photographs (left) and comparison of weight (for each group, fish: *n* = 30), insulin (*n* = 10) and blood glucose (*n* = 10) of fish continuously fed with bacterial fermented feed for 70 days. Asterisks indicate significant differences, based on ANOVA followed by Dunnett test; * *P* < 0.05; ** *P* < 0.01; ns, not significant. **(d)** Survival rate of fish fed with bacterial fermented feed for 70 days after infection with *Aeromonas hydrophila* or *Edwardsiella tarda* (for each group, fish: *n* = 50). Asterisks to the right of the bacterial strain name indicate significant differences, as determined by the log-rank test (\**P* < 0.05; \*\**P* < 0.01). **(e)** The working model for *ckx2* plants improving fish growth. Cytokinins from *ckx2* plants reprograms the soil microbiota, promoting the enrichment of specific microbes in soil. These soil microbes translocate to the fish gut, altering the abundance of keystone genera, which may enhance short-chain fatty acid biosynthesis and degradation of macromolecular carbohydrates to promote fish growth.

To further investigate the effects of bacterial fermented feed on fish growth, a 30-day continuous feeding trial using fermented feed was conducted. On the 30^th^ day, feed fermented by *C. somerae*, *R. lituseburensis*, *P. goldsteinii* and *B. licheniformis* resulted in significant increase in body weight, accompanied with improved metabolic health markers, such as elevated insulin and reduced blood glucose levels (**Fig. 5c**). Moreover, fish fed with fermented feed also showed significant reductions in cumulative mortality rates against *Aeromonas hydrophila* or *Edwardsiella tarda* infections compared to those without feeding fermented feed (10% - 32%, **Fig. 5d**), which indicated that the faster growth and better antimicrobial capacity might be related to the higher contents of acetic, propionic and butyric acids came from *C. somerae*, *R. lituseburensis*, *P. goldsteinii* and *B. licheniformis* fermentation.

In summary, we propose a working model of these processes (**Fig. 5e**). Cytokinins secreted by the *ckx2* plants reprogram the soil microbiota, promoting the enrichment of specific microbes in soil, such as *B. subtilis* and *P. luteus*. These soil microbes translocate to the fish gut, altering the abundance of keystone genera, which may enhance short-chain fatty acid biosynthesis and degradation of macromolecular carbohydrates to promote fish growth and offer antibacterial capacity.

## Discussion

The rice-fish symbiotic eco-system represents a globally significant agroecological practice that enhances food and nutrition security and increases economic profits for millions of smallholder farmers^42^. As fish are the primary source of economic gain in the system, improving fish growth and disease resilience is crucial. Here, we reported that rice mutant with loss function of *Osgn1a* gene could promote grain yield and fish productivity in a rice-fish co-culture system. Compared to WT plants, *ckx2* plants secreted higher levels of cytokinins into the soil, reshaping the soil microbial community and influencing keystone taxa in the fish gut, which in turn improved fish growth and disease resistance. These findings highlight that selecting appropriate rice genotypes is a key strategy for optimizing fish yield within rice-fish symbiotic system. Although the null *ckx2* allele is a rare haplotype, it has been introduced into the elite hybrid rice restorer line Shuhui498 (R498)^43,44^, which could be used in a large-scale rice-fishery co-culture system.

Rice-fish symbiotic ecosystems are a longtime traditional practice, as the productivity of this system is considerably higher than that of rice monoculture due to double harvests ^9,45^. The effects of fish on rice are the most extensively studied aspect of these systems: fish swimming and foraging activities at the base of rice plants help loosen the soil and facilitate the incorporation of feces and feed into the nitrogen cycle within the soil-water environment, thereby lowering the dependence on chemical fertilizers^11,46,47^. Previous study suggests that pollen shed during rice flowering contributes to growth performance by modulating the gut microbiota of Nile tilapia^5^, and similar results are reported in rice-crab and rice-crayfish symbiotic ecosystems^48,49^. Indeed, root-soil-microbiome management is key to the success of regenerative agriculture^50^. Given the relatively primitive immune system of fish compared to that of mammals, their gut microbiota is particularly susceptible to shaping by environmental microbial communities^23,32^. This suggested the possibility that targeted modulation of soil microbiota could help regulate the fish gut microbiome, thereby enhancing growth performance. Herein, *ckx2* plants secreted cytokinins to promote fish growth via reprogramming soil microbiome and subsequently promoting keystone microbes in fish gut in a rice-fish symbiotic ecosystem, illustrating how plant genetic traits can indirectly modulate aquatic animal physiology via belowground microbial shifts.

The rice *OsCKX2* gene has been considered as a star molecule in breeding and genetics since its identification, primarily due to its profound impact on grain yield^39^. *OsCKX2* was first cloned as a major quantitative trait locus *Gn1a*, and cytokinin accumulation in the shoot apical meristem directly promotes the formation of more reproductive organs, thereby significantly increasing the number of grains per panicle^16,18,51^. Moreover, *OsCKX2* gene is associated with increased culm thickness and promotes crown root development in plants, thereby shaping morphogenesis of land plants and enhancing lodging resistance^40,52,53^. Previous studies largely focused on the agronomic benefits of cytokinins, in this study, we showed that *ckx2* plants secreted and accumulated cytokinins into the soil, where they reprogrammed the soil microbes and regulated the abundance of *Bacillus* and *Phaeodactylibacter* which beneficially influence the fish gut microbiota. Our work expanded the functional scope of the *ckx2* plants: from a yield trait regulator to a cross-kingdom communicator within the rice-fish pattern, providing a new strategy for sustainable symbiotic ecosystems through targeted genetic engineering rice trait.

Probiotics have been widely adopted in modern intensive aquaculture practices. The supplementation of specific probiotic microbes, such as *Bacillus* and *Lactobacillus*, has been demonstrated to significantly enhance growth performance and improve resistance against common pathogens in fish^54,55^. Among these, the production of short-chain fatty acids through the microbial fermentation of dietary fiber represents a crucial pathway. Short-chain fatty acids are immediate energy sources for fish^56^, and help lower the gut pH, thereby creating an unfavorable environment for pathogenic bacteria^57,58^. Moreover, short-chain fatty acid, such as acetate acid, serve as signaling molecules that regulate insulin secretion in fish ^59^, subsequently influencing nutrient absorption in fish. Herein, we suggested that fermentation of feeds with key bacterial taxa enhanced short-chain fatty acid levels and improved carbohydrate utilization, thereby accelerating fish growth. Moreover, we successfully isolated a strain of *Romboutsia*, which may be associated with health benefits^60,61^. This strain simultaneously produced multiple short-chain fatty acids during feed fermentation, whereas other bacterial strains used as probiotics typically produce only two or three types, indicating its considerable potential for development as a probiotic.

Taken together, *ckx2* plants secrete cytokinins into soil to reprogram soil microbiome, acting as a sustainable repository of beneficial microbes for fish growth. Future research should focus on elucidating the relative contributions of the three individual cytokinins to soil microbes and expanding the scale of validation trials, thereby promoting the widespread adoption of green aquaculture practices inspired by this study.

## Methods

### Establishment of the rice-fish co-culture system

To investigate the effects of rice plants on fish within a rice-fish co-culture system, an initial experiment was conducted in an indoor artificial climate chamber maintained at 28°C with a 12-hour daily photoperiod. The experimental design comprised two treatment groups: one cultivated with WT and the other with *ckx2* rice plants, each with three replicates. When the rice plants reached a height of 40 cm, the water level in the tanks was increased to 20 cm, and 60 common carp (with an average body length of 8 cm and body weight of 8 g) were introduced into each tank. The fish were fed daily with a ration equivalent to 5% of their body weight. Subsequently, on days 15, 30, 40, 50, 60, and 70, five fish and three soil samples were collected from each tank, frozen, and stored for subsequent analysis.

To validate the impact of *ckx2* rice plants on fish growth, a follow-up experiment was performed in an outdoor paddy field system. The paddy field measured 9 m in length and 6 m in width, containing a central water body area of approximately 24 m². Each pond was stocked with 300 common carp. Rice cultivation practices and the timing of fish stocking were consistent with the indoor experiment. On day 70, 200 fish were collected from each pond for the measurement of body length and weight, alongside the collection of 10 fish and three soil samples.

### Determination and Analysis of Fish Composition

The carp collected on day 70 were dissected to obtain muscle tissue samples. The crude protein, crude fat, and amino acid contents were determined following the methodologies stipulated by the Chinese National Standards GB 5009.5-2016, GB 5009.6-2016, and GB/T 17819, respectively.

The fish gut was resuspended with liquid nitrogen, and then homogenized with methanol (80%). Then centrifuged at 15000 rpm for 15 min to remove the protein. The supernatant was added to derivatization reagent and derivatized at 40□ for 40 min. After that, added to water by well vortexing as the diluted sample. The supernatant (90 μL) was homogenized with 10 μL mixed internal standard solution. Finally, injected into the LC-MS/MS system (AB SCIEX ExionLC^TM^AD) for analysis.

### High throughput sequencing of 16S rRNA gene and bioinformatic analysis

The total DNA of shrimp intestine was extracted by QIAamp PowerFecal DNA Kit (Qiagen, Dusseldorf, Germany) following the manufacturer’s directions. The concentration and purity of total DNA were determined by NanoVuePlus Spectrophotometer (GE Healthcare, Massachusetts, USA). Primer pair 515F (5’-GTGCCAGCMGCCGCGGTAA-3’) and 806R (5’-GGACTACHV GGGTWTCTAAT-3’) were used to amplify the V4 region of 16S rRNA gene, which was modified with a barcode tag containing a random 6-base oligos^62^. Sequencing libraries were generated using TruSeq DNA PCR-Free Sample Preparation Kit (Illumina, San Diego, USA) and the library quantity was determined by Qubit 2.0 Fluorometer (Thermo Scientific, Massachusetts, USA). The 16S rDNA amplicon libraries were sequenced by Illumina Hiseq2500 and HiSeq4000 platforms. The raw data were paired-end reads and their average length was 250 bp.

Raw data quality filtered and the ASV profile information were obtained using dada2 method in QIIME2 software (Version 2022.9) following the instructions^63,64^. Taxonomic assignment was conducted using a SILVA 132 reference data. The relative abundance of each ASV was normalized using the standard of sequence number corresponding to the sample with the least number of sequences.

Alpha diversity, which shows the complexity of species in just one sample through the Shannon index and Chao1 index, was calculated using vegan package in R software (Version 4.10). The beta diversity, based on the Bray-Curtis and Canberra distance was calculated using vegan package in R, which was further visualized to evaluate the species complexity differences of samples by principal coordinate analysis (PCoA). Random forests regression was used to identify the biomarkers of healthy and diseased shrimp and predict the probability of disease using the following parameters with randomForest package in R (cv. fold = 10, step = 0.99, replication=50, ntree = 5000). The contribution of soil microbial community for gut microbiome was evaluated by using Source Tracker 2^65^.

### Isolation of keystone microbes in fish gut

To isolate and purify the keystone microbes in fish gut, thirty common carp were randomly selected. Fish gut was dissected in an anaerobic glove box (COY, USA). Fish content was homogenized with PBS, then plated onto TSB and AGAR medium for cultivation. A total of 500 single colonies were randomly selected and subjected to analysis via a RA-Colony Raman-based colony identification system (Hooke Instruments, China), which was conducted by Guangdong Magigene Biotechnology Co., Ltd. Based on clustering results, representative monoclonal colonies were isolated for further investigation. Bacterial identification was based on MALDI-TOF mass spectrometry detection (Antu, China) and 16S rRNA gene sequencing identification. The utilization of different carbon source substrates was conducted using a VITEK® 2 Compact System (BioMerieux, USA).

### Genome sequencing profile of keystone microbes

Bacterial genome DNA was extracted by QIAamp DNA Mini Kit (Qiagen, USA), and sequencing was performed on both ONT (Nanopore, USA) and Illumina PE 150 platforms (Illumina, USA) by Beijing Biomarker Technologies Co., LTD. The raw data format of the off-machine data from Nanopore sequencing is the binary fast5 format, which contains all the original sequencing signals, with each single read corresponding to an individual fast5 file. After base calling using Guppy (Version 3.2.6), the fast5-formatted data is converted into fastq format for subsequent analysis. Then, after further filtering out low-quality reads and short fragments (length <2000 bp), a final dataset was obtained. For genome assembly, the filtered reads were assembled by Canu software (Version 1.5), and then circlator v1.5.5 was taken to cyclizing assembly genome^66^. For genome component prediction, coding genes prediction was performed by Prodigal (Version 2.6.3), and the GenBlastA (Version 1.0.4) program was used to scan the whole genomes after masking predicted functional genes^67^. For functional annotation, the predicted protein was blast (e-value: 1e-5) against KEGG database (Version 201703).

### Fluorescence-labeled bacteria procedure

Bacterial cultures were harvested at OD□□□ = 0.5, washed once with a buffer solution containing 272 mM sucrose, 1 mM MgCl□, and 7 mM NaH□PO□-Na□HPO□, and finally resuspended in 2 mL of the same buffer. Then, 5 μL of a green fluorescent plasmid (Biosci Bio Ltd.) was added to 90 μL of prepared competent cells, followed by incubation on ice for 5 minutes. After electroporation at 7.5 kV/cm, the cells were plated on chloramphenicol-containing medium. Multiple colonies were subsequently selected for fluorescence observation to confirm the successful introduction of the fluorescent plasmid into the bacteria.

### Feed fermentation and supplement procedure

Bacterial cultures were grown to an OD□□□ of 0.5. Then, 10 g of feed, 20 mL of GAM medium (Hopbio, China), and 50 μL of the bacterial culture were added to a 50 mL centrifuge tube and subjected to anaerobic fermentation for 48 hours. The fermented feed was flash-frozen in liquid nitrogen, and subsequent quantification of short-chain fatty acids and carbohydrate components was performed by a commercial service provider using gas chromatography (Agilent, USA) by Guangdong Magigene Biotechnology Co., Ltd.

Fish were randomly allocated into nine groups, with 100 individuals per group. They were fed seven different types of bacterially fermented feed (each from a single bacterial strain), one type of mixed-bacterial fermented feed, and one control diet. After 30 days of continuous feeding, 10 fish were randomly sampled from each group for the measurement of blood glucose and insulin levels using blood glucose kit and insulin-Elisa kit (Jiancheng, China).

### Fish survival assay under bacterial infection

To investigate the resistance of fish to opportunistic bacterial pathogens, individuals from different experimental groups were separately introduced into 300 L tanks (n = 50 per group) with water temperature maintained at 28°C. Opportunistic pathogen cultures, including *Aeromonas hydrophila* and *Edwardsiella tarda,* were then introduced into the tanks to achieve a final concentration of 10□ CFU/mL. Fish mortality was monitored every two hours over a subsequent one-week period. The Mantel-Cox (log-rank χ2 test) method was subjected to analyze mortality differences between groups.

### Real time quantitative PCR

Total RNA was reverse transcribed into cDNA using a Evo M-MLV RT Kit (Accurate Biology, China). Real-time PCR was performed with SYBR Green Pro Taq HS Mix (Accurate Biology, China) and 500 nM of each primer on a LightCycle 480 System (Roche, Germany). The levels of expression of genes were determined using the 2^−ΔΔCt^ method after normalization to the internal control gene elongation factor 1 alpha (EF1-α). Sequences of primers used in this study are listed in **Table. S2**.

## Data availability

High-throughput sequencing data used in this study was uploaded in NCBI SRA under PRJNA1367677.

## Acknowledgements

This work was financially supported by National Natural Science Foundation of China (42206099 and 32473181), Fundamental Research Funds for the Central Universities (24xkjc023), the earmarked fund for China Agriculture Research System (CARS-48), China-ASEAN Maritime Cooperation Fund, China-ASEAN Center for Joint Research and Promotion of Marine Aquaculture Technology, Moreover, we sincerely appreciate Guangzhou Haizhu National Wetland Park for providing the experimental paddy field for the rice-fish co-culture study.

## Authors contributions

All authors contributed experimental assistance and intellectual input to this study. The original concept was conceived by S. Z, Y. Z, J. H and Y. C. Rice cultivation and Fish farming was conducted by J. L, J. Y, Q. J and S. W. Sample collections were performed by S. Z, R. H, P. H, Z. D, X. T and T. L. High-throughput sequencing and data analyses were performed by S. Z, Z. H, and R. Z. The manuscript was written by S. Z, Y. Z, J. H and Y. C. All authors read and approved the final manuscript.

