## Supplementary Information for "*Gn1a* rice promotes symbiotic fish growth via reprogramming soil microbiome"

### Supplementary Information For:
### *Gn1a* rice promotes symbiotic fish growth via reprogramming soil microbiome

Shen-Zheng Zeng[1,2]†, Yu-Chan Zhang[1]†, Rui-Rui He[1]†, Hui-Yin Pang[1]†, Zhi-Jian Huang[3], Zhi-Xuan Deng[2], Ren-Jun Zhou[2], Xiao-Cheng Tang[2], Tian-Hao Li[2], Juan-Rong Lv[1], Qiao-Juan Huang[1], Jia-Biao Yang[1], Shao-Ping Weng[1], Jian-Guo He[1]*, Yue-Qin Chen[1]*

[1]State Key Laboratory of Biocontrol, Southern Marine Sciences and Engineering Guangdong Laboratory (Zhuhai), School of Life Sciences, Sun Yat-sen University; Guangzhou, 510275, China.

[2]School of Marine Sciences, Sun Yat-sen University; Zhuhai, 519082, China.

[3]School of Agriculture and Biotechnology, Sun Yat-sen University; Shenzheng, 518107, China.

†These authors contributed equally to this work

Dr. Yue-Qin Chen,

**The PDF file includes:**

Figs. S1 to S17

Tables S1 to S2

**Fig. S1. Comparative analysis of muscle components and digestive enzyme of fish.** **(a)** Comparison of muscle components between WT+Fish and *ckx2*+Fish groups (n = 10), including crude fat and protein contents. Asterisks indicate statistical significance (ns: $P > 0.05$, *: $P < 0.05$, **: $P < 0.01$) determined by Student's t-test. **(b)** Comparison of amino acid percentage distribution in fish muscle between two groups (n = 10). Asterisks indicate statistical significance (ns: $P > 0.05$, *: $P < 0.05$, **: $P < 0.01$) determined by t-test. **(c)** Comparison of digestive enzyme between WT+Fish and *ckx2*+Fish groups (n = 10), including amylase, lipase and tryptase. Asterisks indicate statistical significance (ns: $P > 0.05$, *: $P < 0.05$, **: $P < 0.01$) determined by t-test.

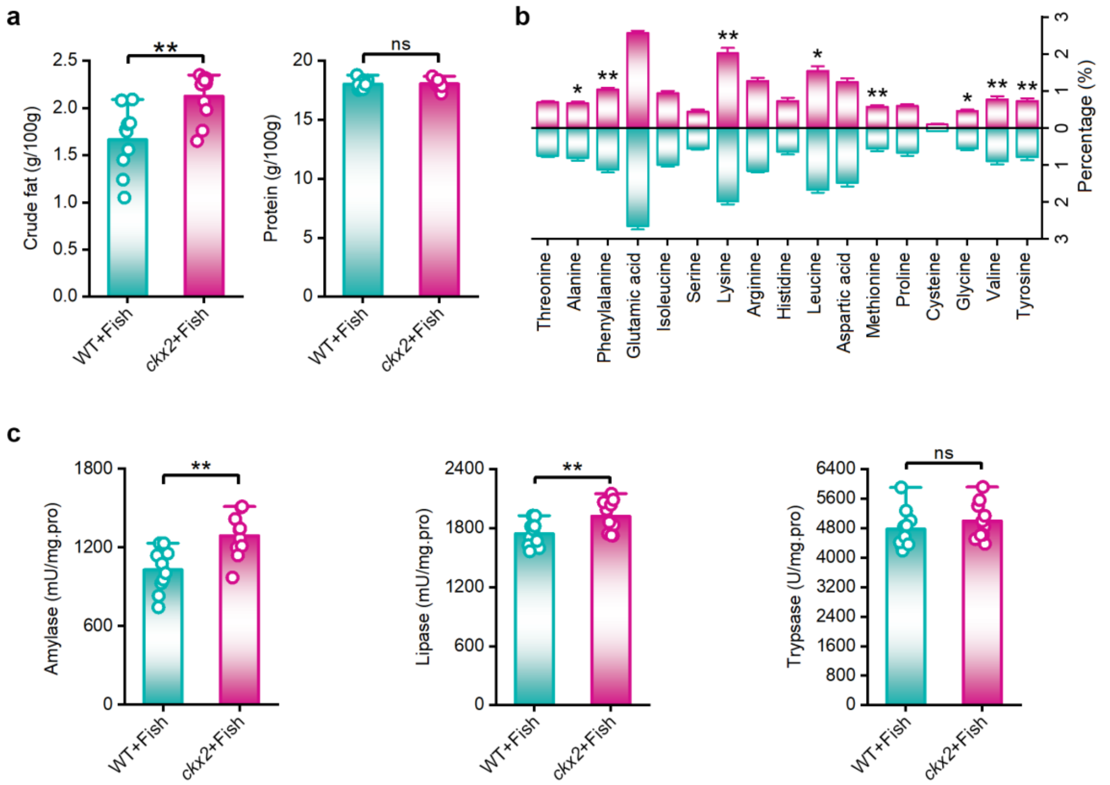

**Fig. S2. Changes gut compounds in response to coculture with *ckx2* plants.**

**(a)** Principal component analysis score plot showed the variance in metabolic profiles between two groups. Green spots represent WT+Fish group, and red spots represent *ckx2*+Fish group. The first two principal components accounted for 36.3% and 24.1% of the total variance, respectively. The PERMANOVA test indicated no significant separation between groups ($P = 0.415$). **(b)** Volcano plot illustrating differential analysis of metabolic features between WT+Fish and *ckx2+Fish* groups. The x-axis represents the log2 fold change, while the y-axis shows the -log10 (*P*-value). Points are color-coded to indicate significantly upregulated (red), downregulated (blue), and non-differentially expressed (gray) features. **(c)** Bubble plot of enriched metabolic pathways comparing *ckx2*+Fish vs. WT+Fish group. The color gradient (red to blue) indicates the -log10 (*P*-value), while bubble size corresponds to the number of differentially expressed metabolites in each pathway.

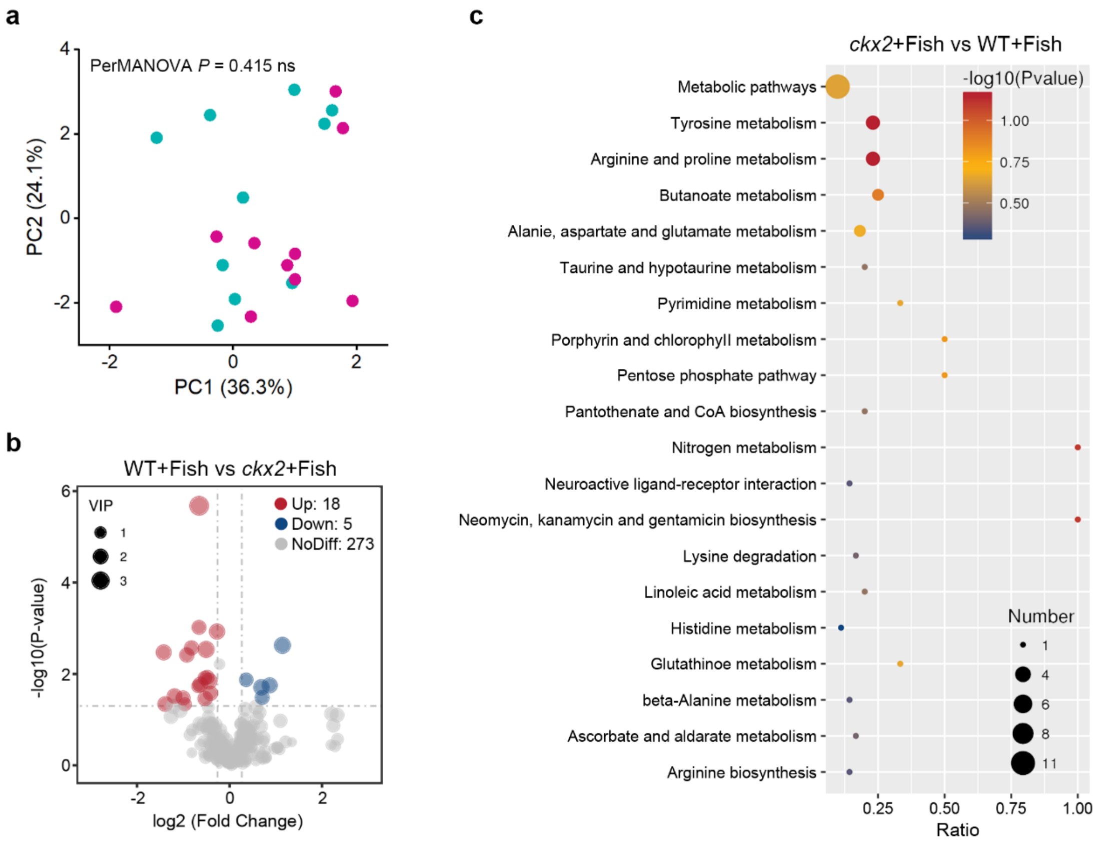

**Fig. S3. Comparative analysis of the expression of fish antimicrobial genes.**

Comparison of the expression of antimicrobial genes between WT+Fish and *ckx2*+Fish groups (n = 10). Asterisks indicate statistical significance (ns: $P > 0.05$, *: $P < 0.05$, **: $P < 0.01$) determined by t-test.

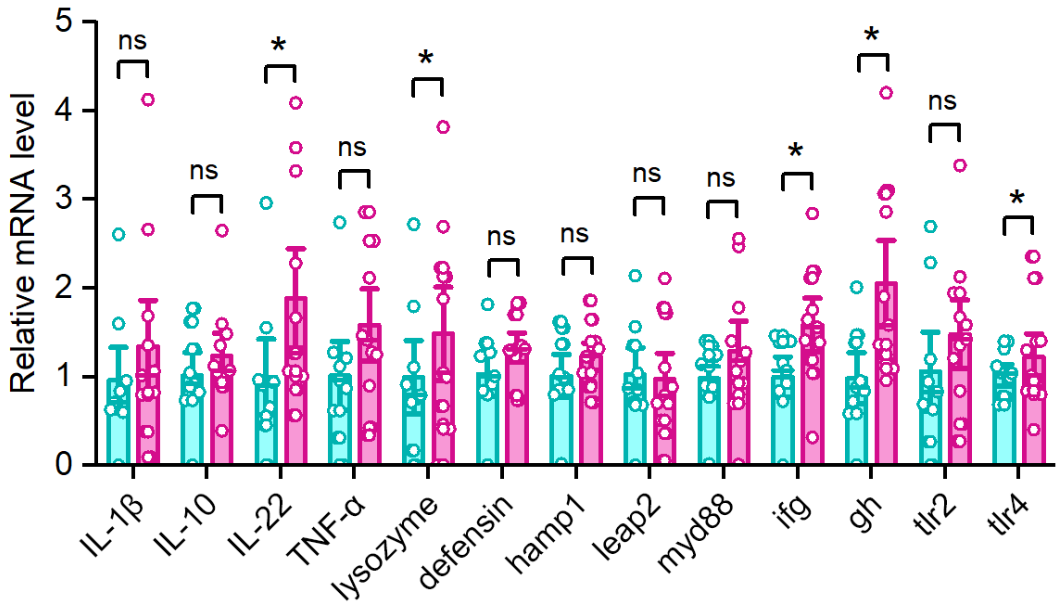

**Fig. S4. Comparative analysis of growth between WT+Fish and *ckx2*+Fish groups of the paddy-pond experiment in 2025 spring.**

Representative photographs (left) and quantification of weight (right) from fish co-cultured with WT or *ckx2* plants at 70 DAC. Scale bars, 6 cm. The asterisk indicates significant differences, based on a t-test; * $P < 0.05$ ($n = 400$).

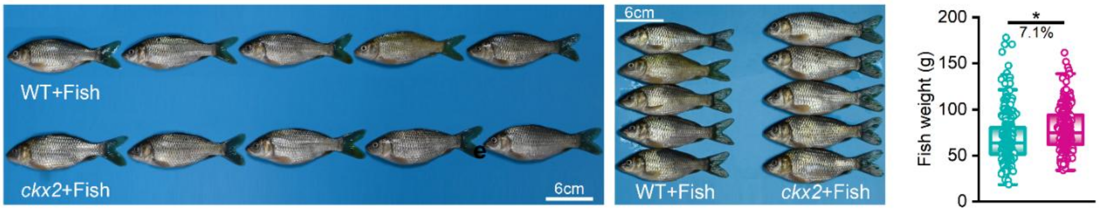

**Fig. S5. The composition in soil and gut microbial community.**

**(a)** The relative abundance of bacterial communities at phylum level in soil. Fusobacteriota, Proteobacteria and Firmicutes were the most abundant phyla. **(b)** The relative abundance of bacterial communities at phylum level in gut. Proteobacteria, Bacteroidota and Acidobacteriota were the most abundant phyla. **(c)** The relative abundance of bacterial communities at phylum level in soil. *Anaeromyxobacter*, *SBR1031* and *Rokubacteriales* were the most abundant genera. **(d)** The relative abundance of bacterial communities at phylum level in gut. *Cetobacterium*, *Aeromonas*, and *Romboutsia* were the most abundant genera.

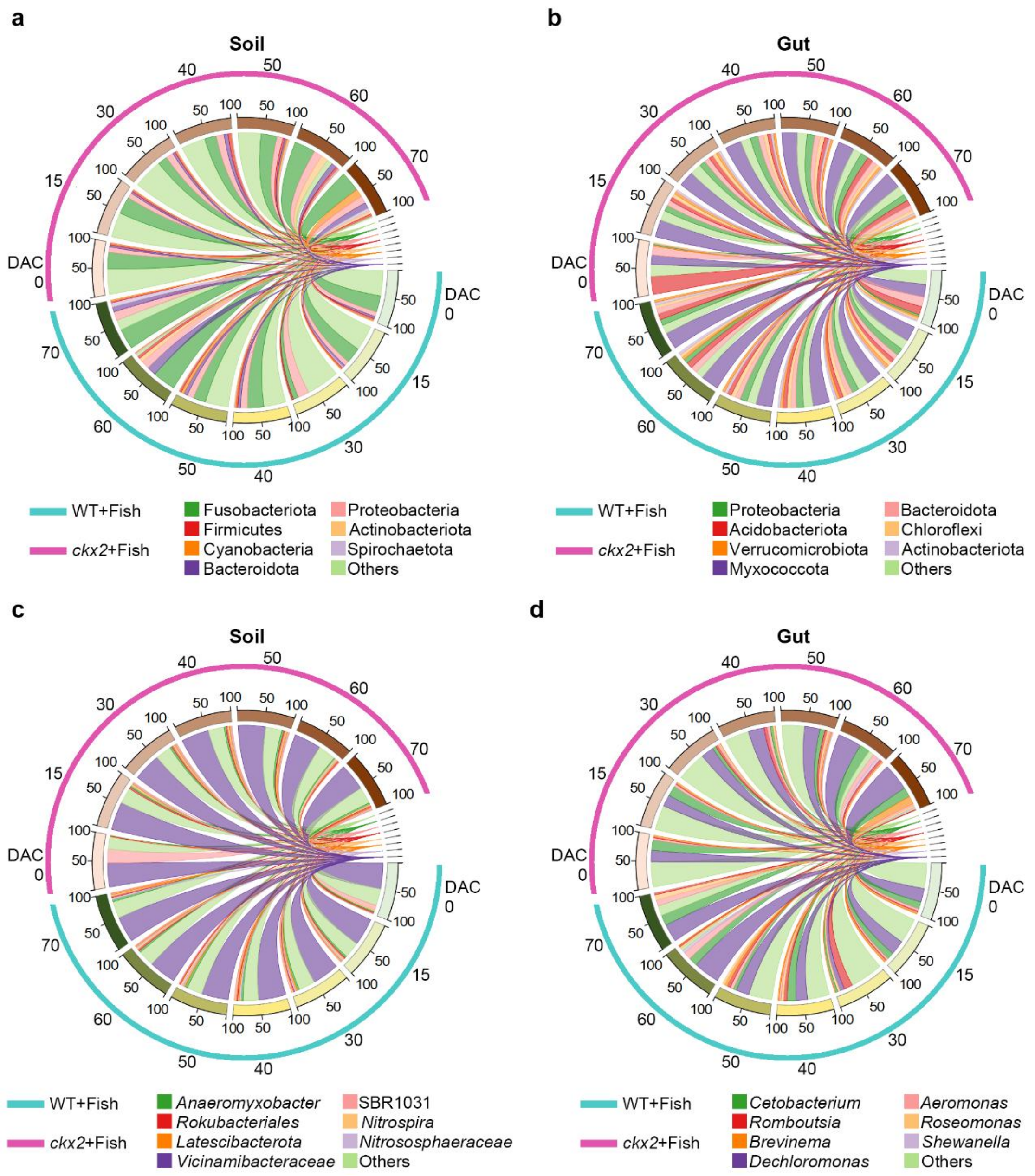

**Fig. S6. Identification of biomarker between WT+Fish and *ckx2*+Fish groups by a random forest model based on genus level.**

**(a)** To find the optimal number of genus-markers for *ckx2*+Fish group, a 10-fold cross-validation on random forest model was conducted. The inflection point where the CV-Error curve began to plateau was selected as the optimal value for biomarkers. **(b)** The classification accuracy of the random forest model was evaluated using the receiver operating characteristic curve (ROC), where a higher area under curve (AUC) value indicated better model performance.

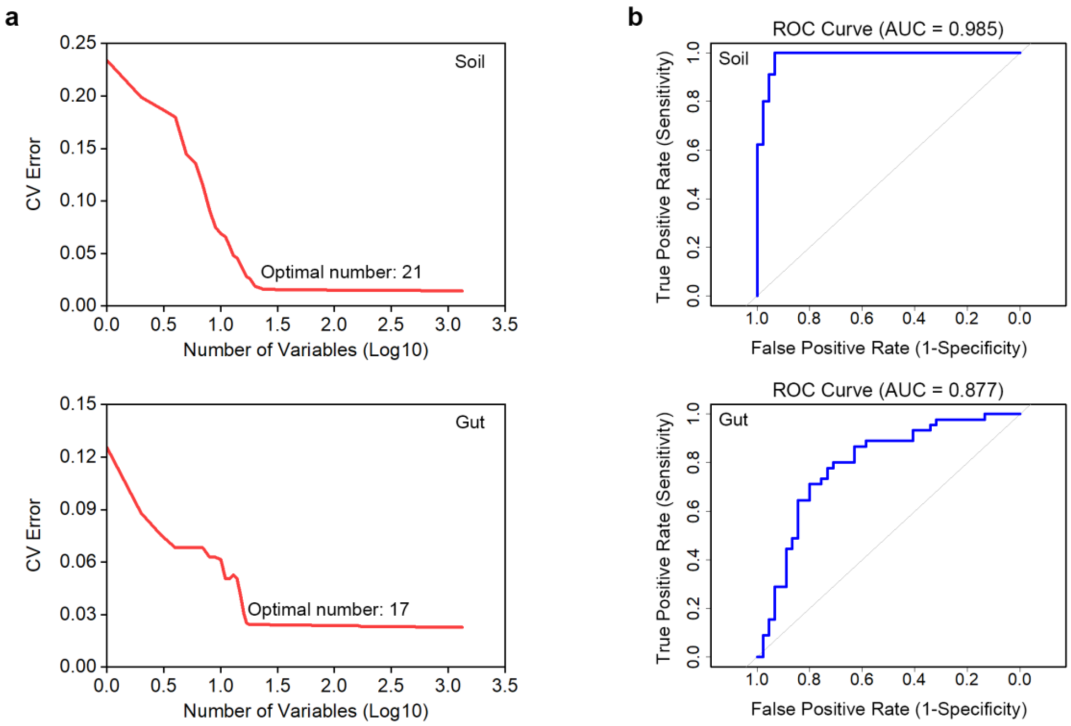

**Fig. S7. Clustering heatmap based on the microbial biomarkers from random forest model.**

**(a)** Heatmaps of soil samples in WT+Fish and *ckx2*+Fish groups based on the 21 biomarker genera. Since divergence in beta diversity appeared at DAC 30, clustering tree showed a high classification accuracy (60.0% - 77.8%) at five timepoints based on the biomarkers. **(b)** Heatmaps of gut samples in WT+Fish and *ckx2*+Fish groups based on the 17 biomarker genera. Since divergence in beta diversity appeared at DAC 50, clustering tree showed a high classification accuracy (60.0% - 80.0%) at three timepoints based on the biomarkers.

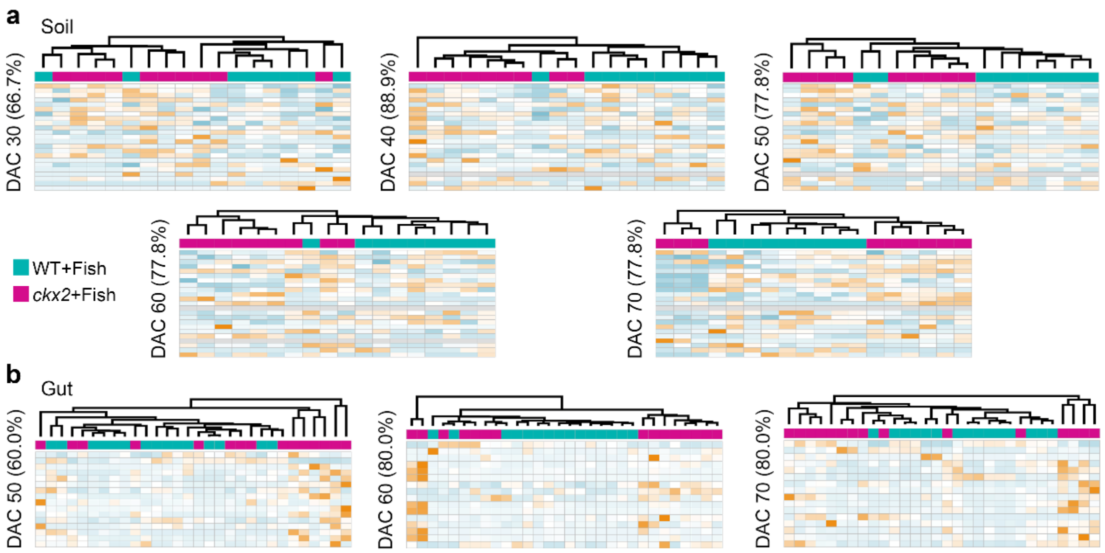

**Fig. S8. Comparison of complexity and stability of species-interaction network between WT+Fish and *ckx2*+Fish groups at different timepoints.**

**(a)** Microbial co-occurrence networks at different time points. Node size corresponded to relative abundance, with larger nodes indicating higher abundance. Edges represented significant co-occurrence relationships between taxa. **(b)** Bar plots showing network topological indexes: average connectivity (left) and average path length (right) for WT+Fish and *ckx2*+Fish networks across time points.

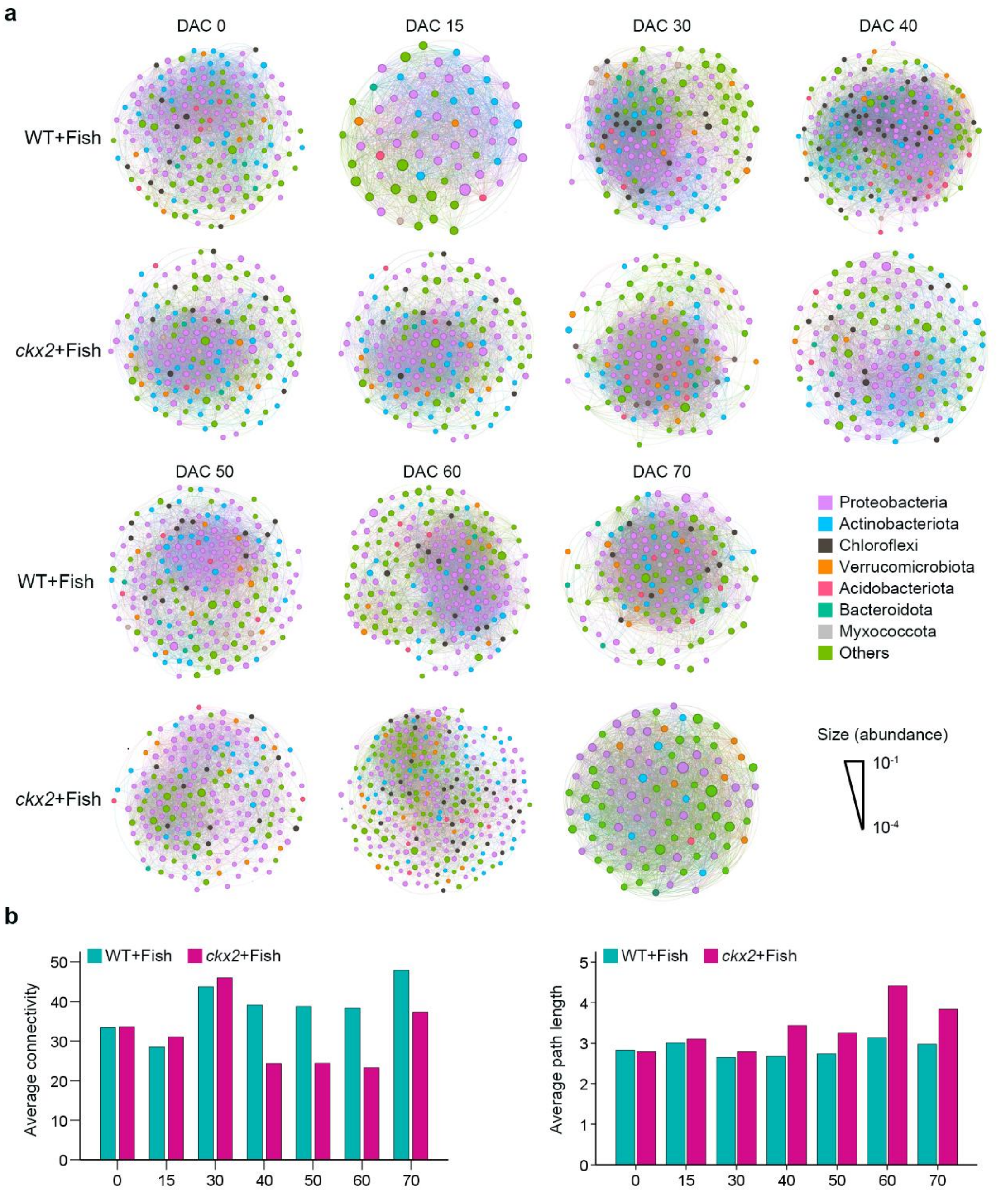

**Fig. S9. Identification of hub microbes in fish gut based on microbial co-occurrence networks**

**(a)** Microbial co-occurrence network based on 210 fish gut microbiomes. Node size corresponded to relative abundance, with larger nodes indicating higher abundance. **(b)** The ASV hubs information of the microbial co-occurrence network. ASVs were sorted by the node degree.

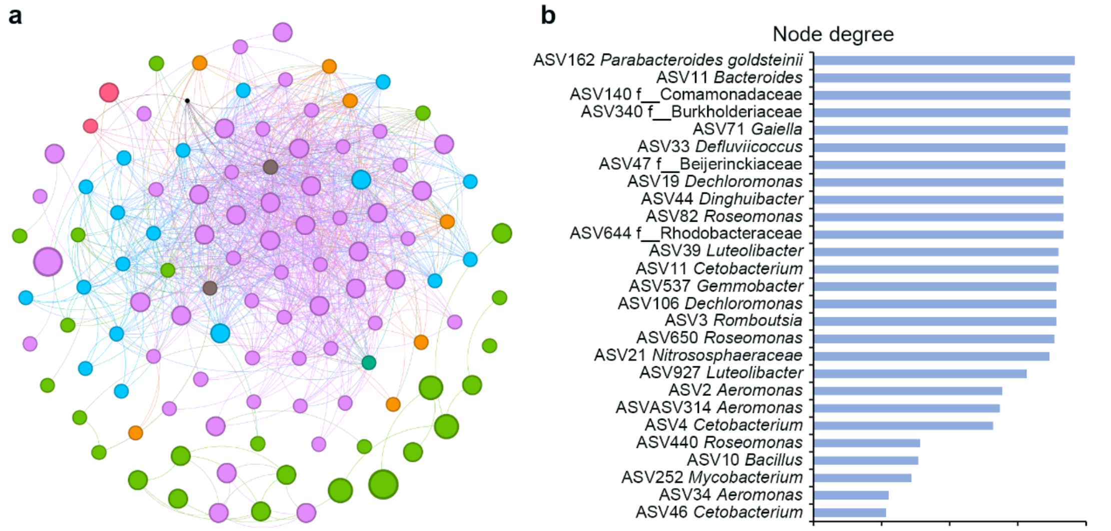

**Fig. S10. The microbial functional difference in soil and gut microbial communities between WT+Fish and *ckx2* groups.**

**(a)** Heatmap depicted the relative abundance levels of various metabolic pathways in soil samples at different sampling timepoints. The color gradient from blue to red illustrates the abundance normalized by Z score. **(b)** Heatmap depicted the relative abundance levels of various metabolic pathways in gut samples at different sampling timepoints. The color gradient from blue to red illustrates the abundance normalized by Z score.

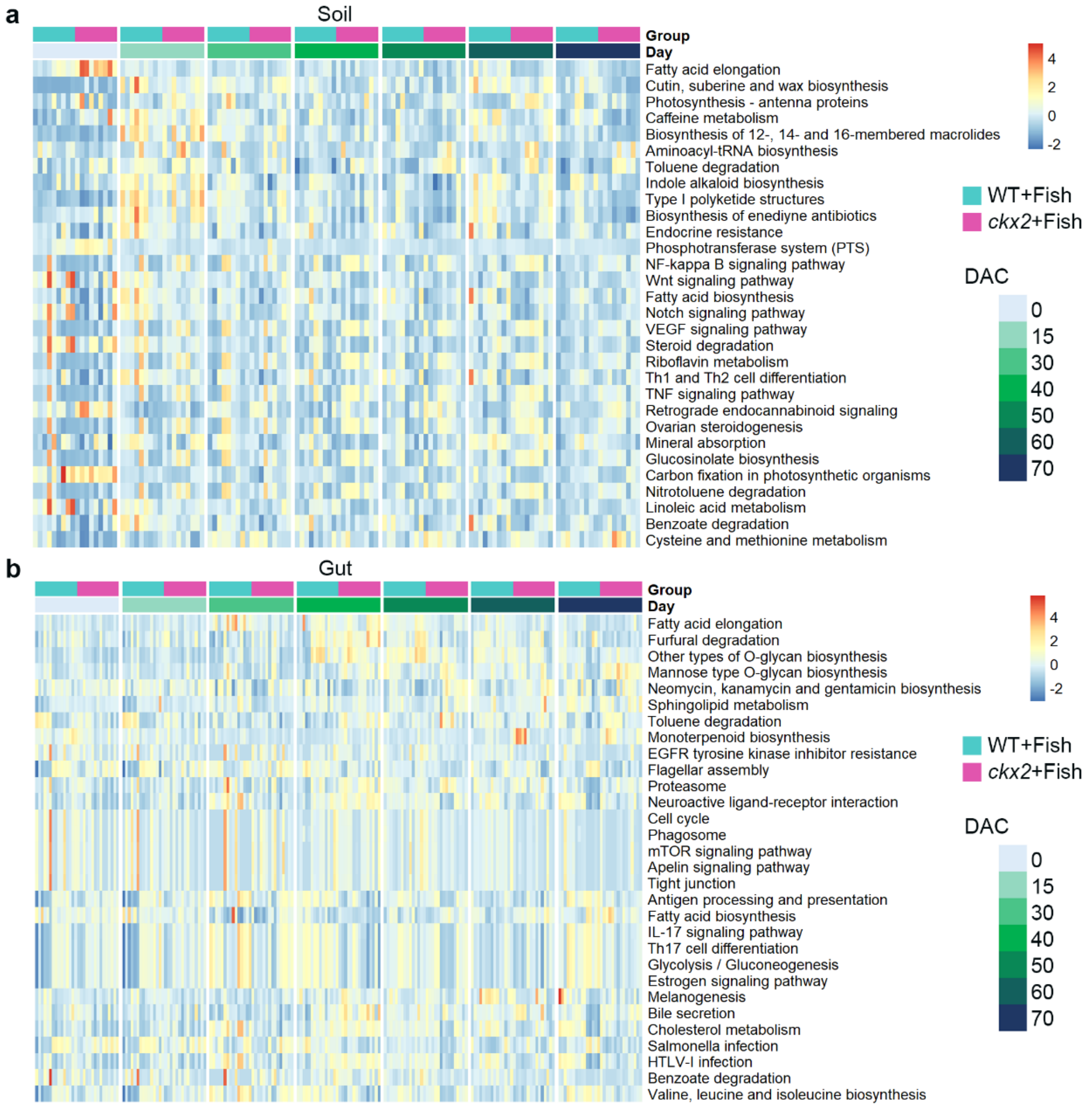

**Fig. S11. Comparison in microbial structure and composition of between control and CK-addition groups.**

**(a)** Differences in microbial structure (left) and composition of soil microbiome between groups ($n$ = 10). Principal coordinate analysis based on Bray-Curtis distance was applied to evaluate the difference in microbial structure between WT+Fish and WT+6-BA+Fish groups. Significant difference ($P$ = 0.001**) in microbial structure between groups was assessed by PerMANOVA. Green triangles indicate the WT+Fish group, and purple triangles indicate the WT+6-BA+Fish group. The abundance of soil biomarkers showed distinctive pattern between two groups. **(b)** Differences in microbial structure (left) and composition of gut microbiome between groups ($n$ = 20). Principal coordinate analysis based on Bray-Curtis distance was applied to evaluate the difference in microbial structure between WT+Fish and WT+6-BA+Fish groups. Significant difference ($P$ = 0.001**) in microbial structure between groups was assessed by PerMANOVA. Green triangles indicate the WT+Fish group, and purple triangles indicate the WT+6-BA+Fish group. The abundance of soil biomarkers showed distinctive pattern between two groups.

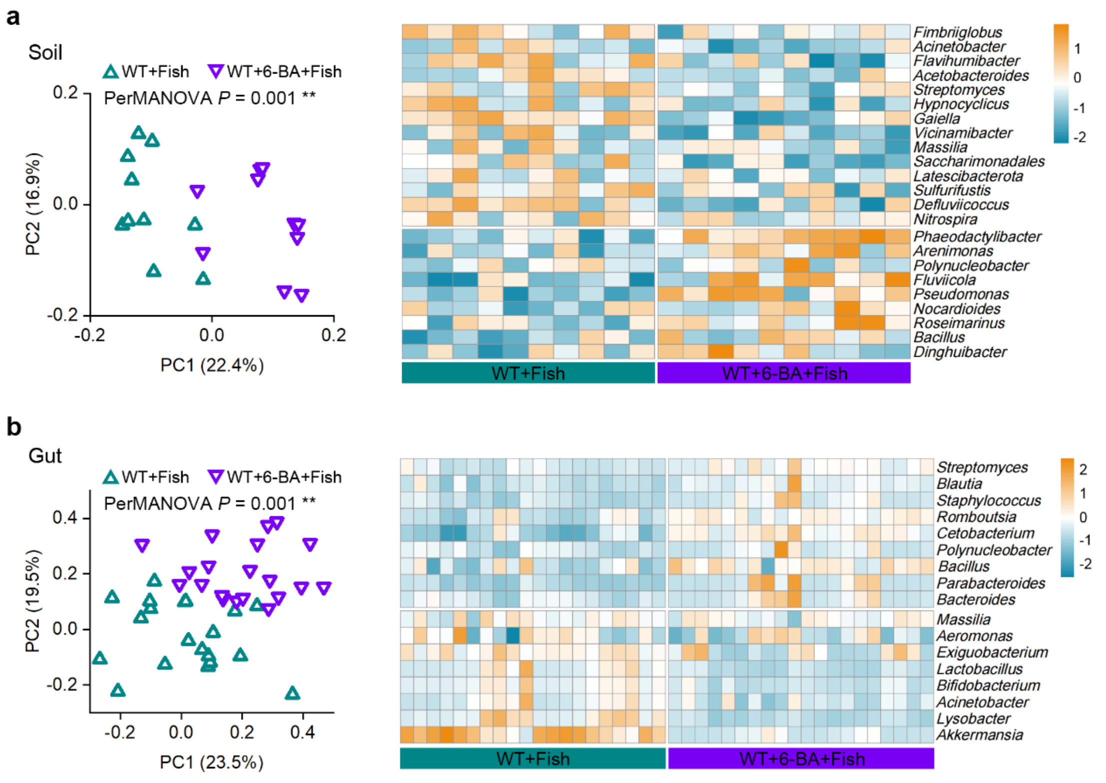

**Fig. S12. Comparison in microbial structure and composition of soil microbiome between control and cytokinin-addition groups.**

**(a)** Principal component analysis plots illustrated the variation in microbial community structures between groups ($n$ = 6). The $P$ value ($P$ = 0.001) suggests significant separation between groups, as determined by PerMANOVA. **(b)** Heatmap of the relative abundance of 21 soil biomarkers between groups. The abundance was normalized by Z-score in the heatmap.

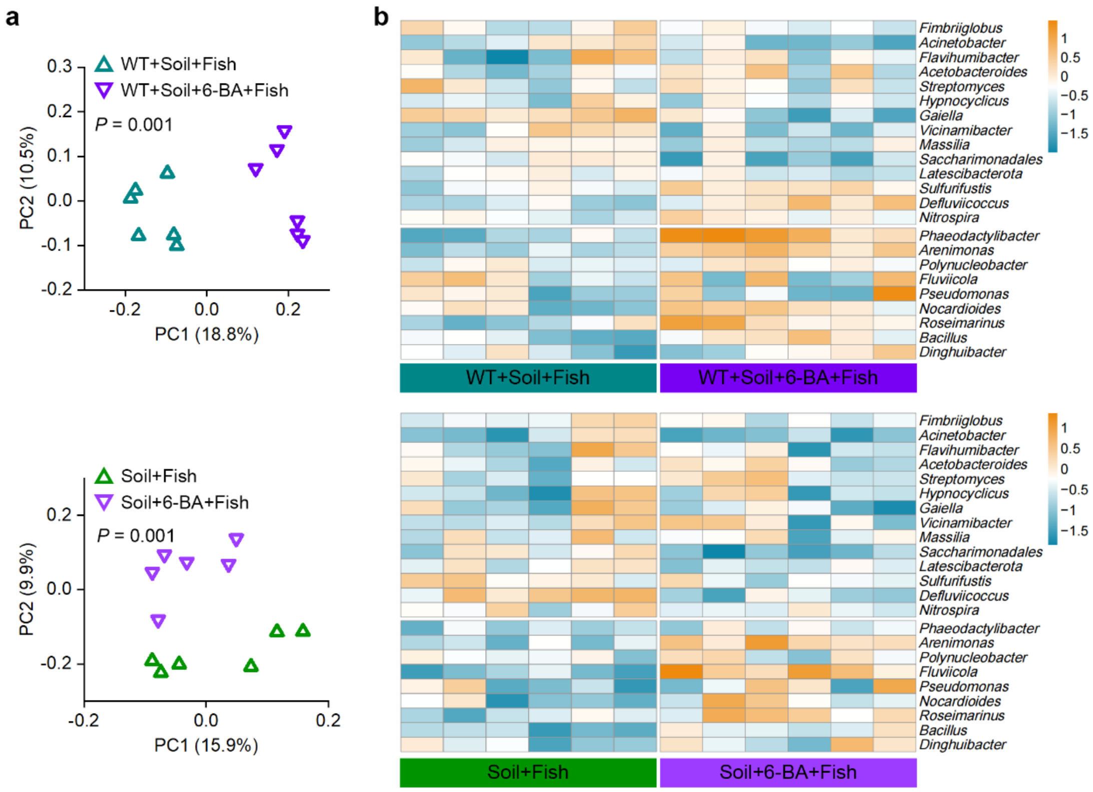

**Fig. S13. Comparison in microbial structure and composition of gut microbiome between control and cytokinin-addition groups.**

**(a)** Principal component analysis plots illustrated the variation in microbial community structures between groups ($n$ = 10). The $P$ value suggests significant separation between groups, as determined by PerMANOVA. **(b)** Heatmap of the relative abundance of 17 gut biomarkers between groups. The abundance was normalized by Z-score in the heatmap.

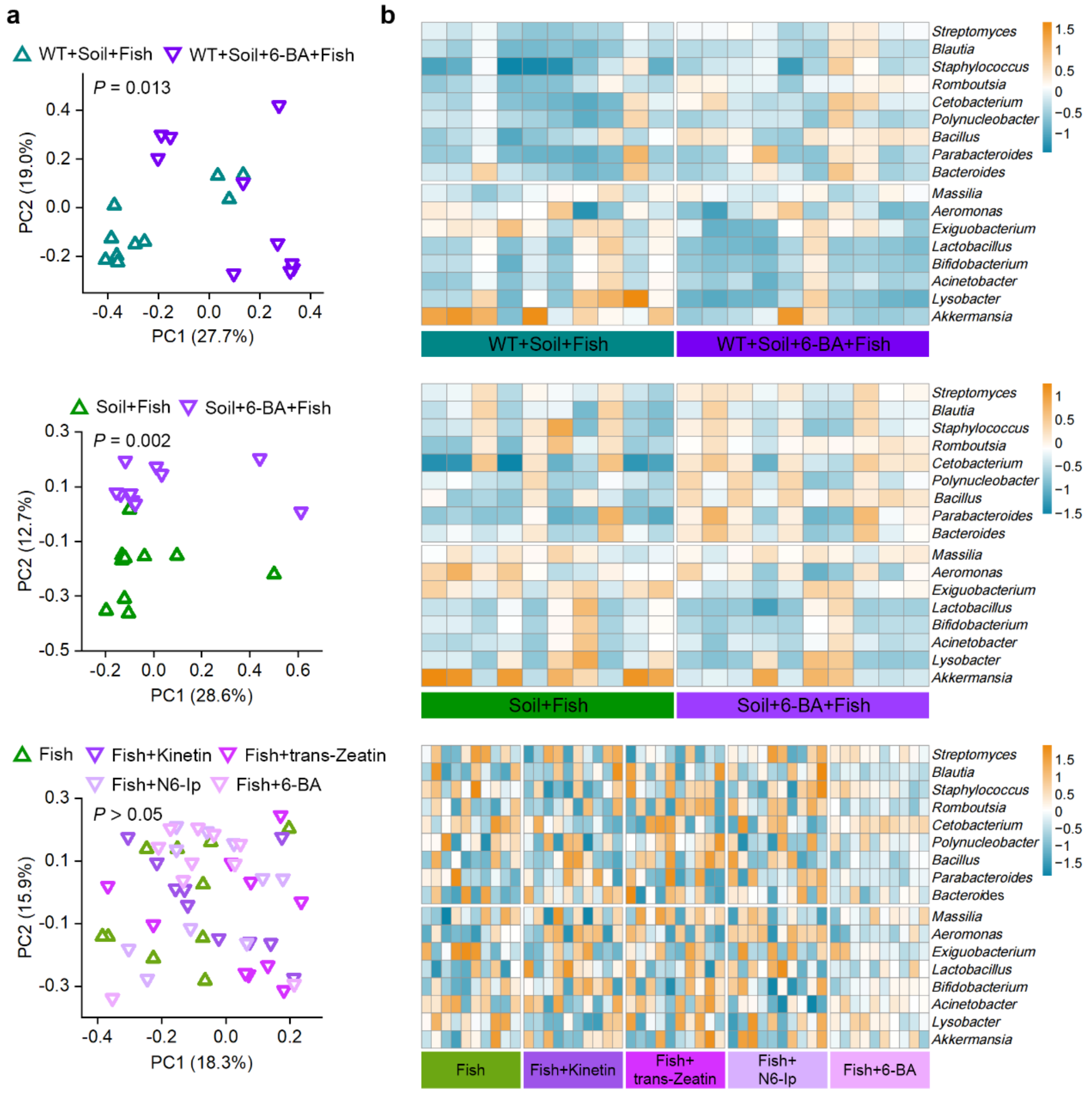

**Fig. S14. Venn diagrams illustrating the overlap of microbial taxa between soil and gut microbiome across timepoints.**

The numbers within the overlapping and non-overlapping regions of the circles indicated the percentage of total abundance of shared or unique ASVs between the soil and gut microbiome.

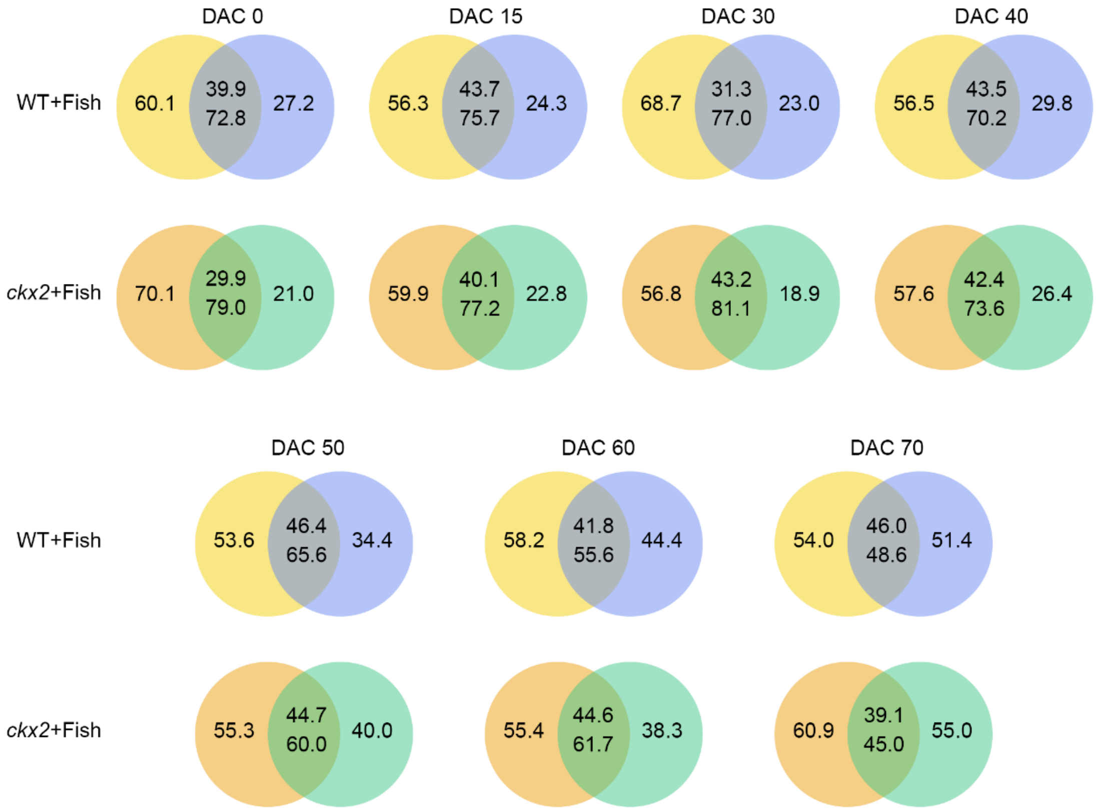

**Fig. S15. The correlation between soil and gut microbiota.**

**(a)** Procrustes correlation is based on the PCoA ordination of the soil microbiota and gut microbiome matrices to calculate the association between two data matrices. Bar chat of Procrustes residual. The *P* value (*P* = 0.031) suggested a significant relation between soil and gut microbiome. The residual value indicated the correlation value between the two data sets. **(b)** Source tracker analysis of contributions of soil microbial community at last time point and gut succession to gut microbiome at next time point, showing a gradually decreasing trend for soil contribution over time.

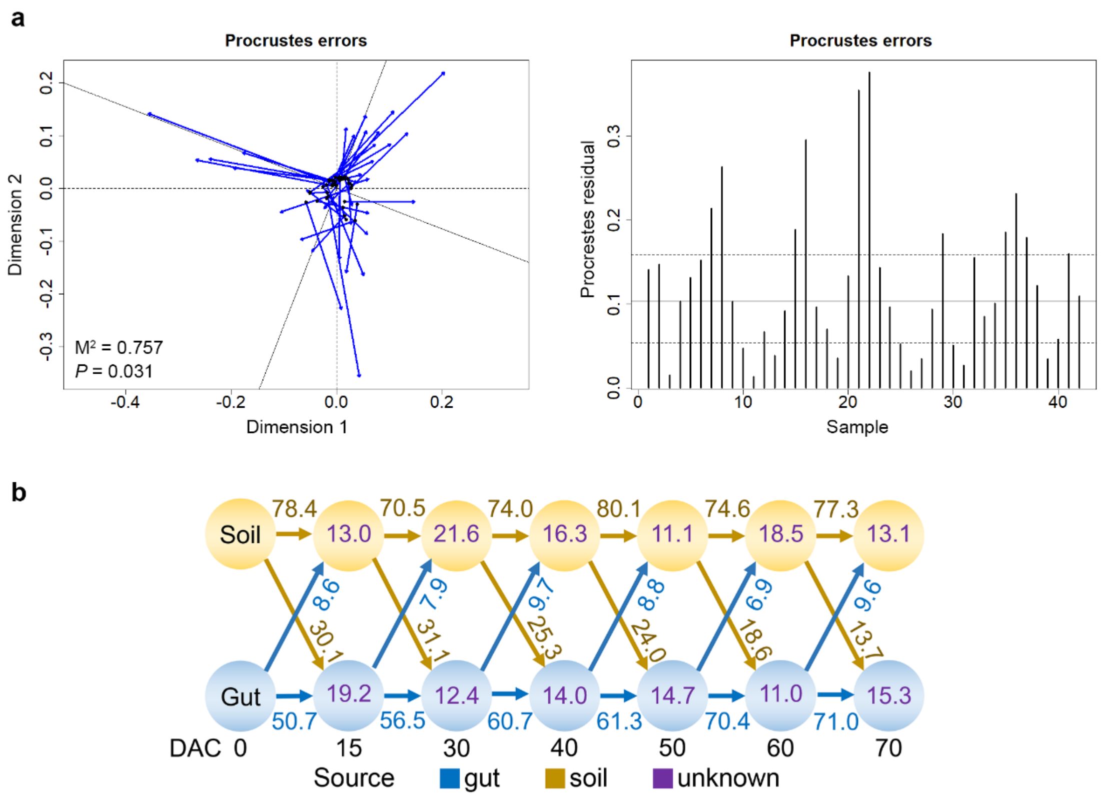

**Fig. S16. Schematic representation of sfGFP plasmid and fluorescent visualization of bacterially transformed colonies.**

**(a)** Circular map of the plasmid pE182-sfGFP plasmid. The plasmid was introduced into bacteria via electroporation. **(b)** Fluorescent images of bacterial colonies expressing green fluorescent protein.

a

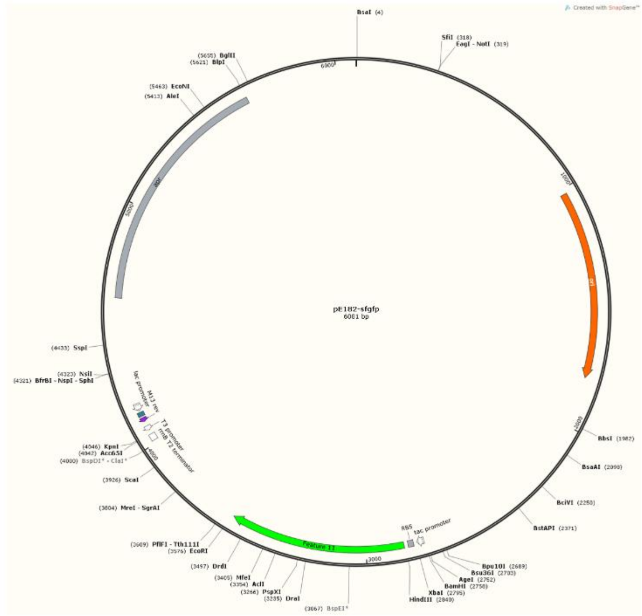

b

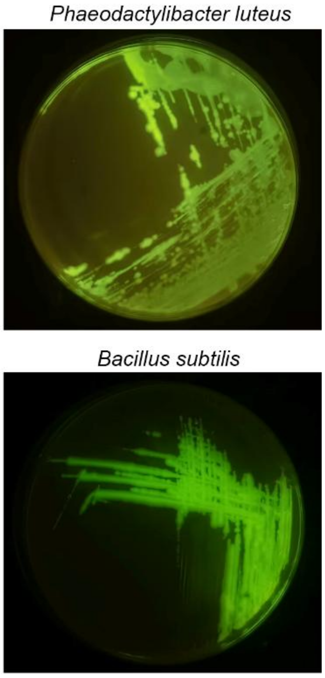

**Fig. S17. Comparative genomic analysis and functional annotation of the seven keystone strains in fish gut.**

Circular genome maps of seven bacterial strains. Each map displayed the genomic architecture providing a visual comparison of genome organization across these strains.

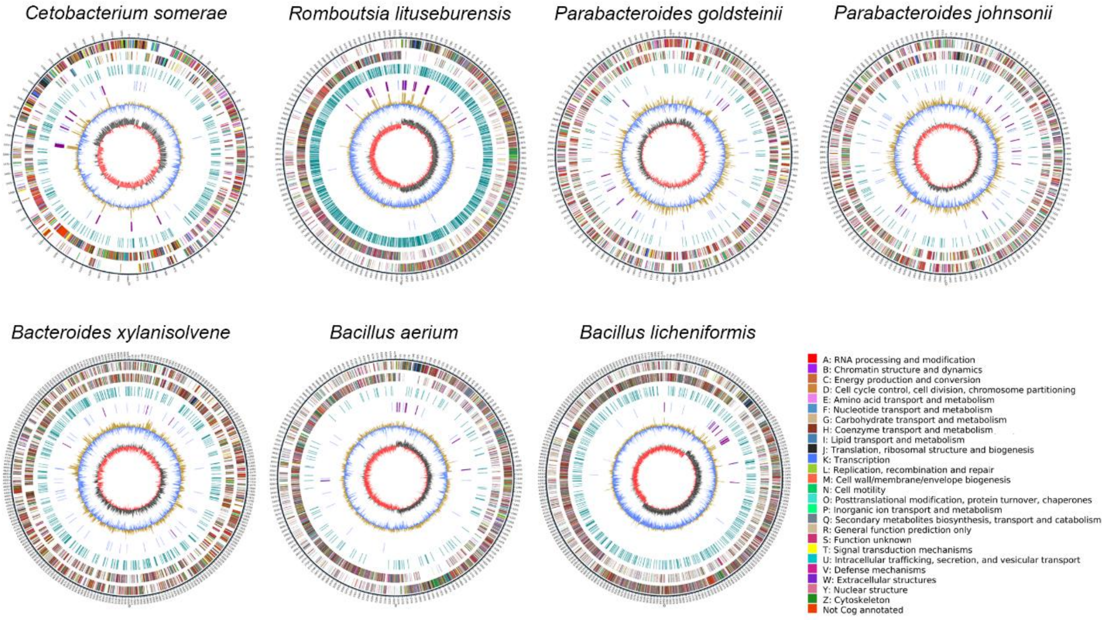

**Table S1. Summary of genome assembly information of the seven keystone microbes.**

| Species | Scaffold Length(bp) | Scaffold Number | Geneset Number | Gene Length (bp) |
|---|---|---|---|---|
| *Cetobacterium somerae* | 2,739,084 | 1 | 2,499 | 2,488,998 |
| *Romboutsia lituseburensis* | 3,883,911 | 2 | 3,556 | 3,290,229 |
| *Parabacteroides goldsteinii* | 4,150,536 | 4 | 3,327 | 3,647,067 |
| *Parabacteroides johnsonii* | 3,741,735 | 1 | 2,946 | 3,284,604 |
| *Bacteroides xylanisolvens* | 6,139,123 | 2 | 4,930 | 5,536,500 |
| *Bacillus aerium* | 4,379,392 | 1 | 4,504 | 3,833,604 |
| *Bacillus licheniformis* | 6,082,999 | 3 | 6,265 | 5,076,780 |

**Table S2. Primer sequences used for real-time PCR for antimicrobial genes.**

| Primer | Sequence (5'-3') |
|---|---|
| beta-actin-F | GCTCTCTTCCAGCCTTCCTTC |
| beta-actin-r | CGGATGTCCACACCACACTT |
| IL-1β-F | ACCAGCTGGATTTGTCAGAAG |
| IL-1β-R | ACATACTGAATTGAACTTTG |
| IL-10-F | CGCCAGCATAAAGAACTCGT |
| IL-10-R | TGCCAAATACTGCTCGATGT |
| IL-22-F | GAAGATCTGCTGCCTCCACGCCA |
| IL-22-R | GCAGAAGTCCTGCAGGTACGTG |
| TNF-α-F | GATGGCAGCCTTGGAAGTGAC |
| TNF-α-R | TCAGAACAATCAGGAAGGAGGAA |
| hamp1-F | TGGAGAGTGAGGCACACCAGGAG |
| hamp1-R | TGCCAGGGGATTGGTTTG |
| lys-F | CAGGTGGAAAGAACAAGTGCA |
| lys-R | ACATCTTACGCCCCTTACAGT |
| def-F | GCAAAGAGAATGAGGCTGTGT |
| def-R | CACAGCACAAAAATCCCTTGC |
| hamp1-F | TGGAGAGTGAGGCACACCAGGAG |
| hamp1-R | TGCCAGGGGATTGGTTTG |
| ifg-F | CCTGGACAAAGCCACTCTTC |
| ifg-R | CTCTCAGCCATTCGCCTTAC |
| gh-F | GCTGGTTAGTTTGTTGGTAAAC |
| gh-R | TTTTACTCAGCTGTCTGCGTTC |
| tlr2-F | AGGATGCTGCGATTCGGTAG |
| tlr2-R | ATGGGACGAGCTGCTTTCAA |
| tlr4-F | GAACTGAATGGGAATAACTGG |
| tlr4-R | TTCTGACTCGCAATAACCAC |
| myd88-F | AGTTTGCACTCAGCCTTTGC |
| myd88-R | GAAAAGATCGGGGCAGTGCG |
